# The variability in potential biomarkers for cochlear synaptopathy after recreational noise exposure

**DOI:** 10.1101/2021.01.17.427007

**Authors:** Tine Vande Maele, Sarineh Keshishzadeh, Nele De Poortere, Ingeborg Dhooge, Hannah Keppler, Sarah Verhulst

## Abstract

**Purpose:** Speech-in-noise tests and suprathreshold auditory evoked potentials are promising biomarkers to diagnose cochlear synaptopathy (CS) in humans. This study investigated whether these biomarkers changed after recreational noise exposure.

**Method:** The baseline auditory status of 19 normal hearing young adults was analyzed using questionnaires, pure-tone audiometry, speech audiometry and auditory evoked potentials. Nineteen subjects attended a music festival and completed the same tests again at day one, day three and day five after the music festival.

**Results:** No significant relations were found between lifetime noise-exposure history and the hearing tests. Changes in biomarkers from the first session to the follow-up sessions were non-significant, except for speech audiometry, that showed a significant learning effect (performance improvement).

**Conclusions:** Despite the individual variability in pre-festival biomarkers, we did not observe changes related to the noise-exposure dose caused by the attended event. This can indicate the absence of noise-exposure-driven cochlear synaptopathy in the study cohort, or reflect that biomarkers were not sensitive enough to detect mild CS. Future research should include a more diverse study cohort, dosimetry and results from test-retest reliability studies to provide more insight into the relationship between recreational noise-exposure and cochlear synaptopathy.

## INTRODUCTION

Noise exposure in young adults mainly takes place during leisure time activities such as watching movies or plays, visiting nightclubs, attending music festivals or concerts, and listening to personal music players. In this population, concerts and music festivals are the most intense activities in terms of loudness (Degeest, Clays, et al., 2017; Keppler, Dhooge, et al., 2015b; Petrescu, 2008; Smith et al., 2000). Sound level measurements during such events confirm that noise levels can exceed 100 dBA (Derebery et al., 2012; Mercier et al., 2003; Ryberg, 2009; Yassi et al., 1993). Despite the potential risk of noise-induced hearing loss (NIHL), the majority of young adults (73.8%) do not wear hearing protection devices (HPDs) when attending concerts (Degeest, Keppler, et al., 2017) or performances at music festivals (86%) (Mercier et al., 2003). After attending these music venues, 36-86% of young adults experience hearing related symptoms (HRS) such as temporary hearing loss, dullness or tinnitus (Keppler, Dhooge, et al., 2015b; Mercier et al., 2003; Smith et al., 2000).

The reduction of hearing sensitivity following noise exposure has extensively been investigated in animal and human studies (Bramhall et al., 2019). The incidence of NIHL depends on many factors, amongst others the noise characteristics: the intensity, frequency and duration of the exposure (Clark, 1991). NIHL can cause permanent threshold shifts (PTS) due to mechanical damage of the cochlear hair cells or supporting cells and metabolic changes that cause cell degeneration and cell death (Wang et al., 2002; Yamane et al., 1995). The reduction of hearing sensitivity after noise exposure can also be reversible. Mechanical damage of outer-hair-cell (OHC) stereocilia, reversible changes in supporting cells, or a swelling of auditory-nerve-fiber (ANF) terminals (Wang et al., 2002) can result in a temporary threshold shift (TTS) which recovers within minutes or hours (Emmerich et al., 2002; Trzaskowski et al., 2014) to weeks (Le Prell et al., 2012; Ryan et al., 2016). Research investigating the pathophysiology of TTS in rodents indicates that excessive glutamate release excitotoxicity is the source of ANF terminal swelling following noise-exposure (Hakuba et al., 2000; Puel et al., 1998). Recent studies have shown that TTS may not be as benign as previously thought, since it can be accompanied by permanent deficits at the synapse level where type-I ANF terminals connect to the inner hair cells (IHCs). Over the past years, this synapse loss has been named cochlear synaptopathy (CS) and has been linked to noise-exposure and aging in mice (Furman et al., 2013; Kujawa & Liberman, 2009; Sergeyenko et al., 2013).

A single IHC contains 10 to 30 presynaptic ribbon-like structures surrounded with glutamate vesicles. Each ribbon connects with a postsynaptic glutamate receptor patch at the ANF terminal side (Liberman et al., 1990). ANFs with low and medium spontaneous firing rates have higher thresholds and connect at the modiolar side of the IHC. They can be distinguished from ANFs with high spontaneous firing rates and lower thresholds, that connect to the pillar side of the IHC (Liberman, 1982). This kind of synapse is unique for the cochlea and retina, and allows fast synaptic vesicle fusion with glutamate release, and fast recovery. Besides encoding a wide dynamic range of sound intensities, the synaptic efficiency enables phase-locking to the stimulus, i.e. action potentials follow the temporal precision of the stimulus fine structure and its envelope. The phase-locked spikes are further encoded in the subcortical and cortical pathway and are important for supra-threshold listening tasks, e.g., speech intelligibility and sound localization (Henry et al., 2016; Safieddine et al., 2012).

Damage to the IHC synapses due to noise exposure, as seen in CS, can occur immediately after noise exposure and can precede a slow degeneration of spiral ganglion cells (Lin et al., 2011). Synapses that connect to ANFs with low spontaneous rates were shown to be most vulnerable, whereas ANFs with higher spontaneous rates are more robust to noise damage (Furman et al., 2013). This susceptibility of low-spontaneous-rate ANFs, combined with the distribution pattern wherein multiple ANFs couple to a single IHC, could be the reason why auditory thresholds recover or remain unaffected by permanent synaptic damage. CS could hence be a possible explanation for the pathophysiology of hidden hearing loss, where normal auditory thresholds are accompanied by supra-threshold hearing deficits such as decreased speech intelligibility in noise or elevated amplitude modulation (AM) detection thresholds (Bharadwaj et al., 2015; Bharadwaj et al., 2014; Smith et al., 2019). In this regard, some human studies were able to relate elevated AM-detection threshold and compromised speech intelligibility to noise exposure history: Bharadwaj et al. (2015) showed that young normal-hearing listeners with less exposure history to recreational noise had significantly lower AM-detection thresholds compared those who reported more frequent attendance in events associated with loud sound exposure. Liberman et al. (2016) reported that normal-hearing listeners with music training background had lower word recognition scores in presence of noise than those without music training background. However, there are other studies which did not find a correlation between the noise-exposure history of the study participants and their respective speech intelligibility performance (Fulbright et al., 2017; Grinn et al., 2017; Grose et al., 2017; Guest et al., 2018). It has also been argued that CS may play a role in the origin of symptoms such as tinnitus or hyperacusis (Liberman & Kujawa, 2017; Schaette & McAlpine, 2011).

Animal studies and computational auditory modeling studies have attempted to define biomarkers for diagnosing CS noninvasively and have shown the potential of auditory evoked potentials (AEPs). More specifically, the auditory brainstem response (ABR) and envelope following response (EFR) are promising tests, given that ABR and EFR-based derived metrics relate directly to the number of histologically-verified synapse counts in animal models (Kujawa & Liberman, 2009; Möhrle et al., 2016; Parthasarathy & Kujawa, 2018; Shaheen et al., 2015). Supra-threshold ABRs are generated by the synchronous firing of ANFs and brainstem neurons in response to the rapid onset of a brief stimulus (Hall, 2007). In rodents with CS, a reduced amplitude of wave I or reduced wave I/V amplitude ratio was observed, which captures the decreased integrity of the peripheral auditory nerve (Furman et al., 2013; Kujawa & Liberman, 2009, 2015; Lin et al., 2011). The EFR is elicited by continuous, amplitude-modulated signals and the strength of the response to the stimulus envelope modulation frequency reflects temporal coding precision in the subcortical neurons (Bidelman et al., 2015; Purcell et al., 2004). Animals or models with CS showed reduced EFR magnitude, which reflects a deficit in temporal coding fidelity of the auditory system (Bharadwaj et al., 2014; Parthasarathy et al., 2018; Shaheen et al., 2015; Vasilkov & Verhulst, 2019; Verhulst et al., 2018), even when hearing sensitivity recovered or remained normal.

Relating these AEP biomarkers to CS in living humans is far more difficult. Comorbidity with OHC-loss and inter-subject variability of AEP measures in humans complicate the use of these methods. Moreover, CS diagnosis through direct synapse counts is not possible in living humans and only few human temporal bone studies have been conducted in relation to CS (Makary et al., 2011; Viana et al., 2015). Therefore, other methods have to be used to evaluate how AEP markers could relate to CS in living humans. Some studies were able to relate ABR waveform amplitudes and EFR magnitude to noise-exposure history (Bramhall et al., 2017; Grose et al., 2017; Paul et al., 2017; Skoe & Tufts, 2018), aging (Garrett & Verhulst, 2019; Konrad-Martin et al., 2012; Vasilkov & Verhulst, 2019) or tinnitus (Bramhall et al., 2018; Schaette & McAlpine, 2011). Garrett et al. (2020) observed a strong correlation between a rectangularly amplitude modulated (RAM) EFR magnitude and speech intelligibility of young and older normal-hearing and hearing impaired subjects. In this regard, Mepani et al. (2021) found a weak but significant correlation between RAM-EFR magnitude and word recognition scores of normal-hearing listeners. However, other studies did not confirm these findings, e.g. ABR wave I amplitude did not correlate with noise-exposure history in (Fulbright et al., 2017; Guest et al., 2018; Prendergast, Guest, et al., 2017; Spankovich, Le Prell, et al., 2017) or no significant relation between ABR wave-I amplitude, wave I/V ratio, EFR magnitude and noise-exposure history was seen in (Guest et al., 2018). Longitudinal studies in humans, where AEP-measurements are used to evaluate auditory-nerve integrity before and after noise-exposure, are scarce. One study reported unchanged compound action potential (AP) amplitudes in electrocochleography, the analogue of ABR wave I amplitude, in relation to recreational noise exposure (Grinn et al., 2017).

This study investigates the relationship between recreational noise-exposure and markers of CS and OHC damage to understand whether this exposure results in TTS, CS and/or speech in noise deficits. By adopting a test battery that includes pure tone audiometry (PTA) at conventional and extended high frequencies (EHFs), speech audiometry tests in quiet (SPiQ) and in noise (SPiN) and AEP measurements, we hypothesize that speech intelligibility and AEP measurements may deteriorate due to noise-exposure, even if PTA is not affected or recovered shortly after noise exposure. Multiple speech audiometry conditions, ABR and EFR stimuli were used to study the relation of outcome measures to the individual noise-exposure history (Part I) and to analyze the variability in these measures before and in a period of five days after attending a music festival (Part II).

## METHODS

### PARTICIPANTS

Young adults aged 18 to 25 years who planned to attend a music festival during the summer of 2019 were recruited for this study. Volunteers completed a recruiting questionnaire, which was used to exclude subjects with known hearing pathologies, history of ear surgery or tinnitus. Study participation consisted of four measurement sessions: one baseline measurement session (session 1) between 1 to 3 days before the start of the festival and 3 follow-up sessions: 1 day, 3 days and 5 days after the end of the music festival. Further on, these follow-up sessions will respectively be referred to as session 2, session 3 and session 4. Subjects were asked not to expose themselves to noise other than that of the festival during the follow-up period and not to use party drugs. Participants were free to use hearing protection.

Twenty subjects participated in the first session, 8 males and 12 females (mean age 21.5 years ± 2.24 SD; standard deviation). Of those subjects, the best ear was chosen as the test ear based on tympanometry and PTA. This was the right ear for 7 subjects (4 males) and the left ear for 13 subjects (4 males). One male subject (right testing ear) dropped out of the study for the last measurement session, resulting in 19 participating subjects for the follow-up analysis in Part II, with mean age of 21.6 years (SD 2.27). At every test session, subjects completed a test battery consisting of questionnaires, PTA, SPiQ and SPiN tests, and AEP measurements. The test protocol had a duration of maximum 3 hours per session (including breaks and information) and tests were conducted in the same order for every subject and at every session.

This study was approved by the ethical committee of the Ghent University Hospital (Belgium) and was performed following the statement of ethical principles of the Declaration of Helsinki. All subjects were informed about the testing procedures and signed an informed consent. Subjects received a financial compensation for their participation.

### QUESTIONNAIRES

The questionnaire of the baseline session consisted of four parts and was completed online or during the first session. The first part consisted of sociodemographic questions, questions regarding present HRS, illness, recent drug use and noise exposure. Finally, the general use of hearing protection devices and the type of hearing protection was queried. The second part consisted of questions regarding noise-exposure history based on a Dutch translation of Jokitulppo et al. (2006) by Keppler, Dhooge, et al. (2015b). A list of relevant, noisy recreational activities was presented. First, subjects estimated the frequency of noise-exposure caused by each activity and the duration of each exposure in hours. Second, the exposure level (Laeq) was estimated using a relative scale based on communication effort, ranging from 60 dBA (sound level of a normal conversation) to 100 dBA (sound level which makes communication impossible). The use of HPD was asked and converted in the equation as a Laeq-reduction of 10 dB. Finally, for each activity, the total number of years that this activity was present in their lives on the one hand, and for the last three years on the other, was asked. Cumulative lifetime-noise-exposure history (Laeq,life) was calculated using the formula of Jokitulppo et al. (2006). An analogous calculation was made for recent noise-exposure history (Laeq,recent), which represents cumulative noise exposure over the past three years. The third and fourth part respectively consisted of the Youth Attitude to Noise Scale (YANS), (Widen et al., 2006) and Beliefs about Hearing Protection and Hearing Loss (BAHPHL), (Svensson et al., 2004) as modified by Keppler et al. (Keppler, Dhooge, Degeest, et al., 2015; Keppler, Dhooge, et al., 2015a; Keppler & Dhooge, 2010). Both questionnaires use a five-degree Likert Scale ranging from “totally disagree” to “totally agree” to determine the attitudes and beliefs towards noise, hearing protection and hearing loss. A higher score reflects more positive attitudes towards noise, less concerns about hearing loss and less belief of benefit of HPDs.

At sessions 2, 3 and 4 a questionnaire that evaluates HRS, recent drug and alcohol consumption, and between-session noise-exposure was completed. The first follow-up session contained additional questions regarding the festival duration and noise-exposure. For each level of communication effort, the subject was asked to estimate a percentage of exposure time and whether HPDs were used in this listening situation to determine the festival noise-exposure (Laeq,festival) based on a 40-hour-week noise-exposure calculation (Jokitulppo et al., 2006; Keppler, Dhooge, et al., 2015b).

### OTOSCOPY AND TYMPANOMETRY

Otoscopic inspection of the ear canal and tympanic membrane was performed at the baseline session. All test ears had normal otoscopic results. Middle-ear admittance was bilaterally measured at the baseline session, using a GSI TympStar (Grason-Stadler) tympanometer with a 226 Hz, 85 dB sound pressure level (SPL) probe tone. All test ear tympanograms had an ear canal volume between 0.6 and 2.0 mmho, a static acoustic admittance level within a range of 0.3 to 1.7 mmho and a middle-ear pressure between −100 and 100 daPa, and were defined as normal type-A tympanograms.

### PURE-TONE AUDIOMETRY

PTA thresholds were measured while subjects were seated in a double-walled sound-attenuating measurement booth. At the baseline session, both ears were tested to select the test ear. Air-conduction thresholds were measured at conventional octave frequencies 0.125, 0.250, 0.500, 1, 2, 4 and 8 kHz and half-octave frequencies 3 and 6 kHz, and at EHFs of 10, 12.5, 14 and 16 kHz using the modified Hughson-Westlake procedure. An Equinox Interacoustics audiometer was used and stimuli were transmitted using Interacoustics TDH-39 headphones and Sennheiser HAD-200 headphones for conventional frequencies and EHF, respectively. Maximum output levels for extended high frequencies of 10 kHz, 12 kHz, 14 kHz and 16 kHz were 80, 70, 60 and 40 dB HL, respectively. If no threshold was reached at the maximum output level, the threshold was considered as this maximum stimulus level for data analysis. However, this only occurred in one subject at 16 kHz. The ear with better thresholds on conventional frequencies was chosen as the test ear, provided that these thresholds were all 20 dB HL or better, and that this ear had a normal otoscopic evaluation and a type-A tympanogram. Since air-conduction thresholds on conventional frequencies never exceeded 20 dB HL and type-A tympanograms showed normal middle-ear admittance in all test ears, bone conduction measurements were not performed.

### SPEECH INTELLIGIBILITY IN QUIET AND IN NOISE

At each session, SPiQ and SPiN tests were performed at the test ear using the Flemish Matrix sentence test (Luts et al., 2014) in Apex 3 software (Francart et al., 2008). Sentences were presented in a relatively quiet room using a laptop connected to a Fireface UCX soundcard (RME) and HDA-300 (Sennheiser) headphones.

In each session, five randomly chosen experimental test lists were presented in five differently filtered conditions, presented in a random order. By filtering speech stimuli, different auditory mechanisms can be targeted. Low-pass (LP) filtered speech understanding is thought to mainly rely on temporal fine-structure cues, whereas high-pass (HP) filtered speech would mostly rely on envelope-coding of the auditory system. Therefore, the five conditions consisted of two conditions in quiet, where speech was filtered with a zero-phase 256^th^-order finite impulse-response (FIR) LP filter or with a zero-phase 1024^th^-order FIR HP filter with cutoff values of 1500 and 1650 Hz, respectively. Three conditions were presented in a speech-shaped noise with a fixed level of 70 dB SPL: a broadband condition (BB) where no filtering was applied, a LP and a HP condition, where speech as well as noise signals were filtered using the same filter cutoff values as for LP and HP condition in quiet, respectively. Due to the filtering, the SPLs of 0 dB SNR LP and HP conditions were 70 and 53 dB SPL, respectively. During the first session, two training lists were presented in the BB-in-noise condition to minimize the known learning effect of this test (Kollmeier et al., 2015). For all test lists, subjects were asked to repeat the five-word sentences in a closed, forced-choice setting (10 options for each word were given). Speech was presented in an adaptive procedure using a staircase paradigm to determine the speech-reception threshold (SRT) with a minimal step size of 0.1 dB. The mean signal level or mean SNR of the six last reversals was used as the SRT for the SPiQ and SPiN-tests, respectively. Lower SRT-values reflect better speech intelligibility.

### AEP MEASUREMENTS

In each session, AEP measurements were performed with the Universal Smart Box system (Intelligent Hearing Systems) using SEPCAM software. Subjects were seated in a comfortable armchair in a quiet room and were watching a muted video during the recordings. The skin preparation was performed using NuPrep gel and Ambu Neuroline snap electrodes were placed on vertex, nasion and bilateral earlobes. The electrode impedances never exceeded 3 kΩ. For one subject, a mastoid electrode configuration was used due to an immovable earring. All stimuli were presented using ER-2 probes (Etymotic Research). Both ears were covered with earmuffs to minimize noise interaction and all inessential electrical devices were turned off during the measurements. Subjects were instructed to lean their head back and to relax as much as possible.

ABRs were collected using 5000 alternating polarity sweeps of six stimuli types: broadband 80-µs click stimuli and narrowband 500-ms toneburst (TB) stimuli. Slow click rates of 11 Hz as well as fast click rates of 120 Hz were both presented at levels of 80 dB and 100 dB peak equivalent SPL (peSPL). Faster click rates cause minimal neural recovery time and have earlier been proposed for neurodiagnostic purposes (Valderrama et al., 2012). TBs with center frequencies of 1 and 4 kHz, which provide more frequency specific information, were presented at 70 dB SPL with a rate of 20 Hz. The raw signals were stored and processed offline in Matlab R2018b using the custom-made “sepcam2mat” function. Recorded ABR traces were bandpass filtered between 100 and 1500 Hz with an 800^th^ order FIR filter using “filtfilt” Matlab function to remove the filter delay. Twenty millisecond epochs were extracted relative to the stimulus onset and baseline correction was applied by subtracting the mean-value of each epoch. Two hundred epochs, equal number of each polarity, with the highest peak-to-trough values were rejected and the remaining 4800 epochs were averaged (Keshishzadeh et al., 2021). Mean ABR level series, starting from 100 dB-peSPL and down, were plotted to visually identify the peaks based on the mean latency intervals by Table 8-1 in Picton (2011). Accordingly, ABR wave I, III and V peaks and latencies were identified manually and were confirmed by visual inspection of a second audiologist. Figure 1 shows exemplar ABRs to 80 and 100 dB peSPL, 11-Hz clicks. Identified peaks are specified with O (ABR to 80 dB peSPL, 11 Hz clicks) and X (ABR to 100 dB peSPL, 11 Hz clicks). The wave I/V amplitude ratio was defined as the ratio of peak-to-baseline amplitudes of wave I and wave V, respectively.

**Figure 1:**
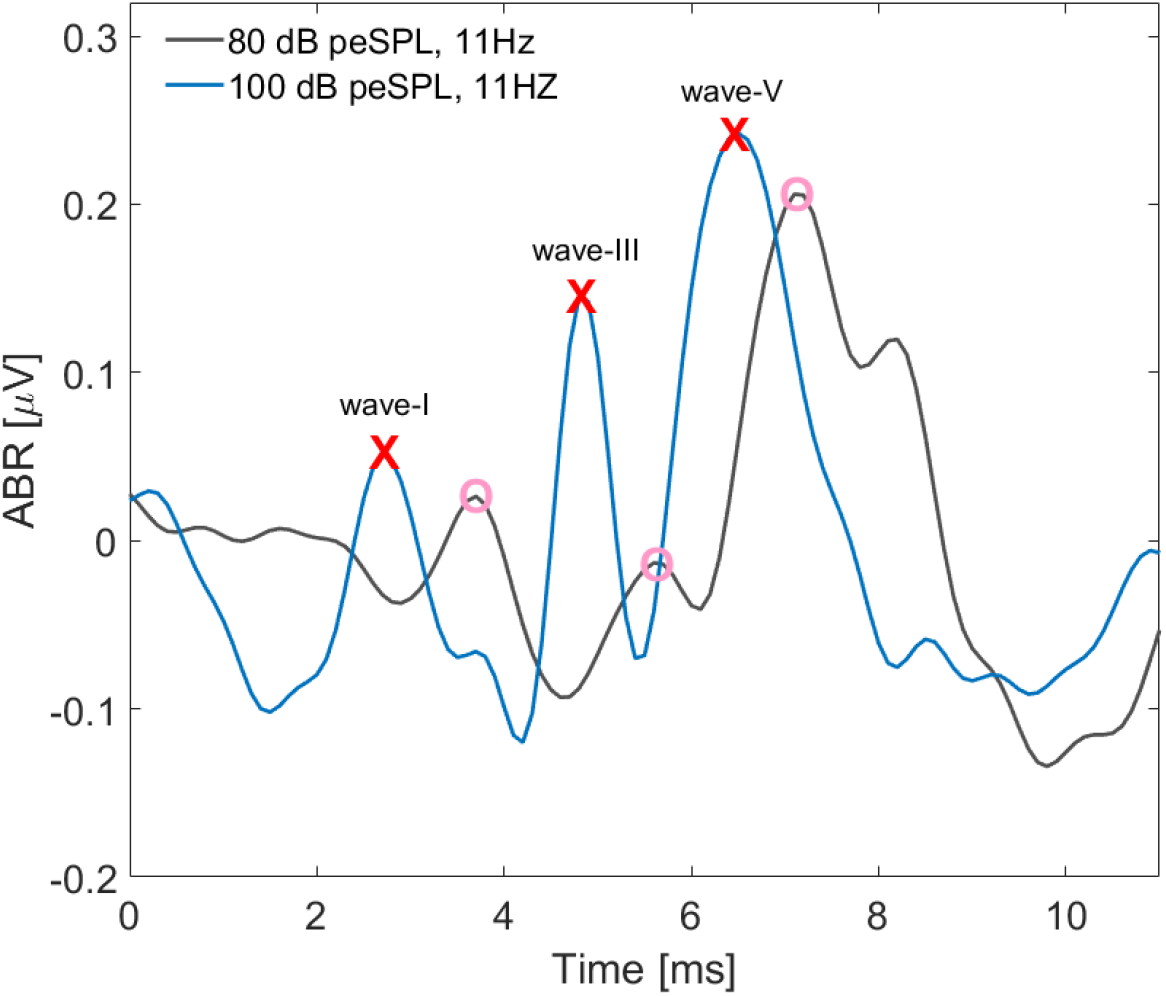
Exemplary ABR waveforms to 80 dB peSPL (dark grey) and 100 dB peSPL (blue), 11 Hz clicks. Respective ABR wave I, III and V peaks are specified by O and X for 80 and 100 dB peSPL click stimuli, respectively.

EFRs were evoked using 1000 sweeps of two stimulus types. Both stimuli consisted of a 4-kHz amplitude-modulated tone and are distinguished by their modulation waveform. Specifically, sinusoidal amplitude modulation (SAM) was applied in the first stimulus and rectangular amplitude modulation (RAM) in the second (25% duty cycle). The latter was adopted from Vasilkov et al. (2021), as this stimulus was found to yield strong EFRs which are sensitive to detect CS in auditory model simulations. Both stimuli had a modulation frequency of 110Hz, a modulation depth of 100%, and were presented at 70dB SPL. Stimuli had a duration of 500 ms and were presented at a rate of 2 Hz.

The EFR processing was performed offline in Matlab R2018b. First, recorded EFRs were bandpass filtered using the same bandpass FIR filter as applied to ABRs, but with low and high cutoff frequencies of 50 Hz and 5000 Hz, respectively. Then, 400-ms epochs were extracted from the 100 to 500-ms time-interval, relative to the stimulus onset and 20 epochs with the highest peak-to-trough values were rejected. The remaining 800 epochs were averaged and the corresponding magnitude spectrum was constructed using the fast Fourier Transform (FFT). Additionally, a bootstrapping approach was adopted in the frequency domain to estimate the noise-floor and variability of the EFR. Exemplar SAM and RAM EFR spectra and corresponding noise-floors are shown in Figure 2A and C, respectively. For detailed explanation of the bootstrapping procedure see Keshishzadeh et al. (2021). EFR strength was defined as the summation of the signal-to-noise spectral magnitude at fundamental frequency and the following three harmonics, i.e. 110, 220, 330 and 440 Hz, if they were above the noise-floor (Vasilkov et al., 2021). Noise-floor corrected SAM and RAM EFRs are shown in Figure 2B and D. Arrows represent the EFR magnitudes at modulation frequency (110 Hz) and existent spectral peaks at corresponding harmonics.

**Figure 2:**
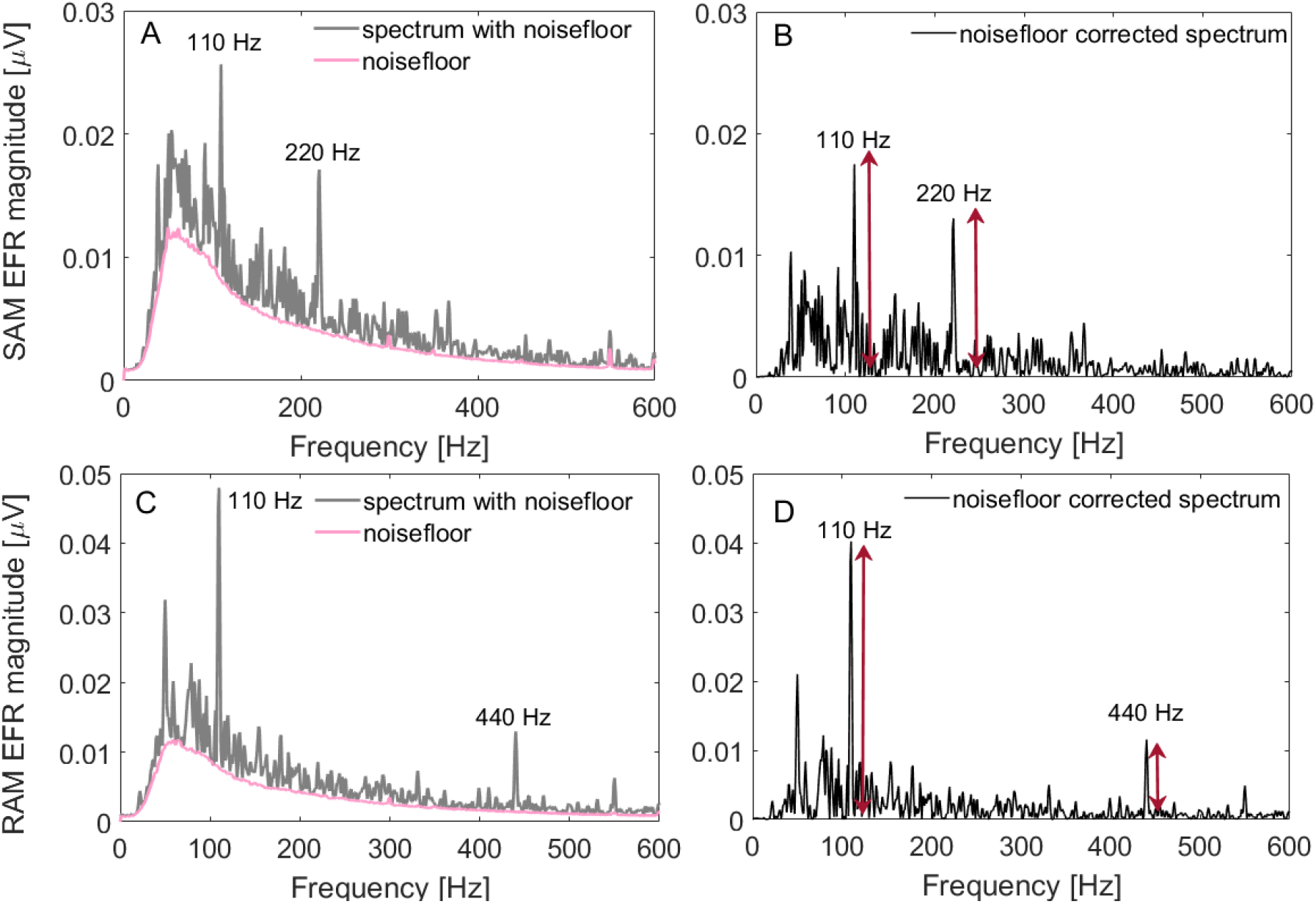
Exemplary EFR spectra. (A) SAM EFR spectrum (grey) and estimated noise-floor (pink). (B) Noise-floor corrected SAM EFR after bootstrapping. (C) RAM EFR spectrum (grey) and estimated noise-floor (pink). (D) Noise-floor corrected RAM EFR after bootstrapping. Arrows in panels (B) and (D) show the magnitudes of EFR at modulation frequency (110 Hz) and harmonics.

### DATA ANALYSIS

Statistical data analysis was performed in two stages using IBM SPSS Statistics 25. Descriptive parameters were calculated and tests of normality – i.e. Shapiro-Wilk test, histograms, Q-Q plots, and box and whisker plots - were applied.

First, the baseline session was analyzed. Hearing-test results of audiometry, speech audiometry SRT, EFR magnitude, ABR amplitudes and latencies of waves I, III and V and I/V amplitude ratio were analyzed in relation to lifetime noise-exposure history. Although Laeq,life was normally distributed as defined by descriptive parameters and the Shapiro-Wilk test, Spearman’s ranked-order correlation coefficients (*ρ*) were reported as outliers and/or not-normally distributed test parameters were present in the data for one or more audiometric frequencies on the one hand, and for one or more test conditions of speech audiometry, EFR or ABR on the other. Two-tailed significance level was reached for correlations when p<0.05, suggesting a monotonic and significant relationship between Laeq,life and the considered hearing-test parameter.

Second, intersession variability was analyzed. As there were outliers and the data was not normally distributed for some sessions or test conditions, the assumptions for using repeated measures analysis of variance (ANOVA) were not met and a non-parametric related-samples Friedman’s two-way ANOVA by ranks was conducted to determine statistically significant changes in all hearing test parameters over sessions. If significance level (p<0.05) was reached, post-hoc analysis of least-square means was performed for intersession differences between session 1 on the one hand and session 2, 3 and 4 on the other. The two-tailed significance level was reached if p<0.017 when applying a Bonferroni correction for multiple comparisons.

For all test results, the shift from the first to the second session was calculated. Non-parametric Mann-Whitney-U tests were conducted to compare those shifts between groups of subjects that reported HRS after the musical event and those who did not. Significance level was reached when p<0.05, suggesting a significant difference between the two groups.

## RESULTS PART I

### QUESTIONNAIRES

Frequencies of Laeq,life and Laeq,recent are shown in Figure 3. Mean Laeq,life was 70.17 dBA (SD 7.83, range 56.46 – 90.16) when corrected for HPD-use. Attending a discotheque or dance party were reported as the loudest activity (m 79.5 dBA, SD 8.87, range 60.0 – 90.0), followed by attending a concert or a festival (m 74.2 dBA, SD 5.07, range 70.0 – 80.0). All but one subject had attended such events before they participated in this study. Noise-exposure history for the last three years, as defined by Laeq,recent, had a mean of 66.29 dBA (SD 7.92, range 54.41 – 84.94). Pearson correlation showed a strong positive, statistically significant correlation between Laeq,recent and Laeq,life (r(20)=0.973, p<0.001). Mean total YANS-score was 2.81 (SD 0.67, range 1.26 – 4.05) and mean total BAHPHL-score was 2.42 (SD 0.47, range 1.50 – 3.38). Twelve subjects (60%) reported commonly wearing HPDs when exposed to loud noises, five subjects (25%) used customized ear plugs.

**Figure 3:**
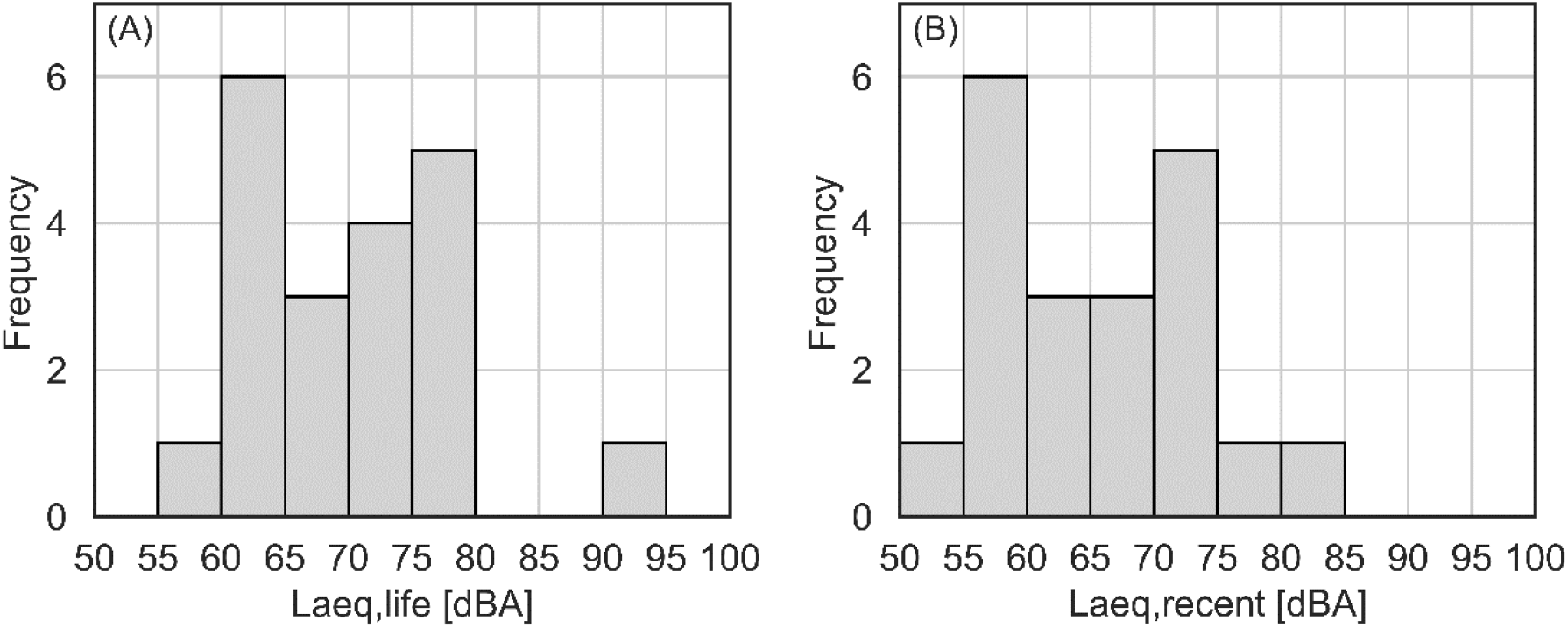
Frequency distribution of Laeq,life (A) and Laeq,recent (B).

### BASELINE HEARING STATUS

Box-plots of the hearing thresholds for conventional frequencies and EHF thresholds of the testing ears are shown in Figure 4. The mean pure-tone average of 3, 4, 6 and 8 kHz (PTA3-8kHz) was 4.88dB HL (SD 2.75, median 5.00, range −1.25 – 8.75) and the mean pure-tone average of 10, 12.5, 14 and 16 kHz (PTA10-16kHz) was 5.94 dB HL (SD 9.59, median 2.50, range −10.00 – 25.00). Neither PTA3-8kHz nor PTA10-16kHz significantly correlated to lifetime noise-exposure history (Laeq,life: p>0.05). Spearman’s correlation coefficients ranged from −0.28 to 0.09. The relation between PTA10-16kHz and Laeq,life and their respective Spearman’s correlation coefficients are shown in Figure 1 and 2 of Supplemental Material.

**Figure 4:**
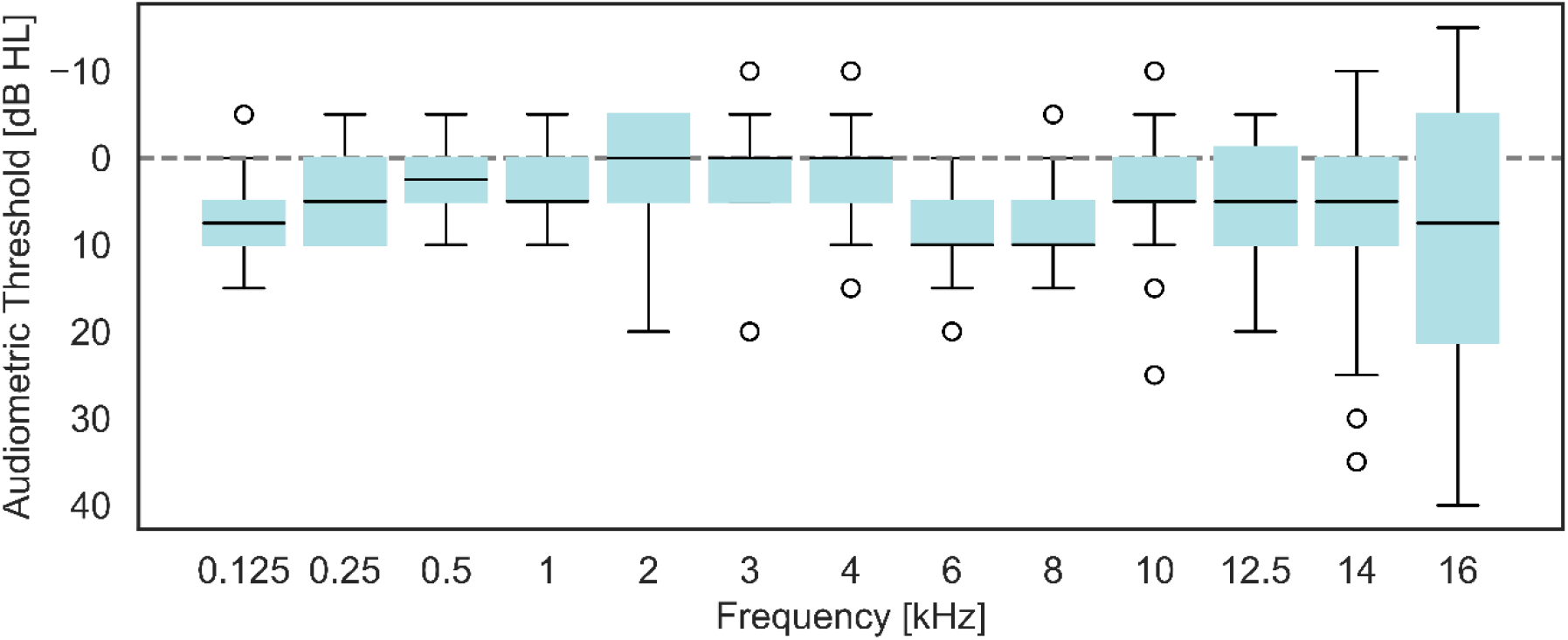
Boxplots of baseline audiometric thresholds.

### BASELINE SPEECH AUDIOMETRY

Figure 5A and B represent the box-plots of SPiQ and SPiN SRT-values, respectively. No monotonic relation was found between SPiQ and Laeq,life, as Spearman’s correlation coefficients were 0.24 (p = 0.30) for the LP filtered speech and −0.20 (p = 0.41) for the HP filtered speech, and both were statistically insignificant. Spearman’s correlations of SPiN to Laeq,life showed weak positive correlation coefficients, ranging from 0.02 to 0.44 with the highest correlation coefficient for the LP condition. However, none of the correlation coefficients were statistically significant (p>0.05). The relation between Laeq,life and speech intelligibility SRT values of different conditions can be found in Figure 3 and 4 of Supplemental Material.

**Figure 5:**
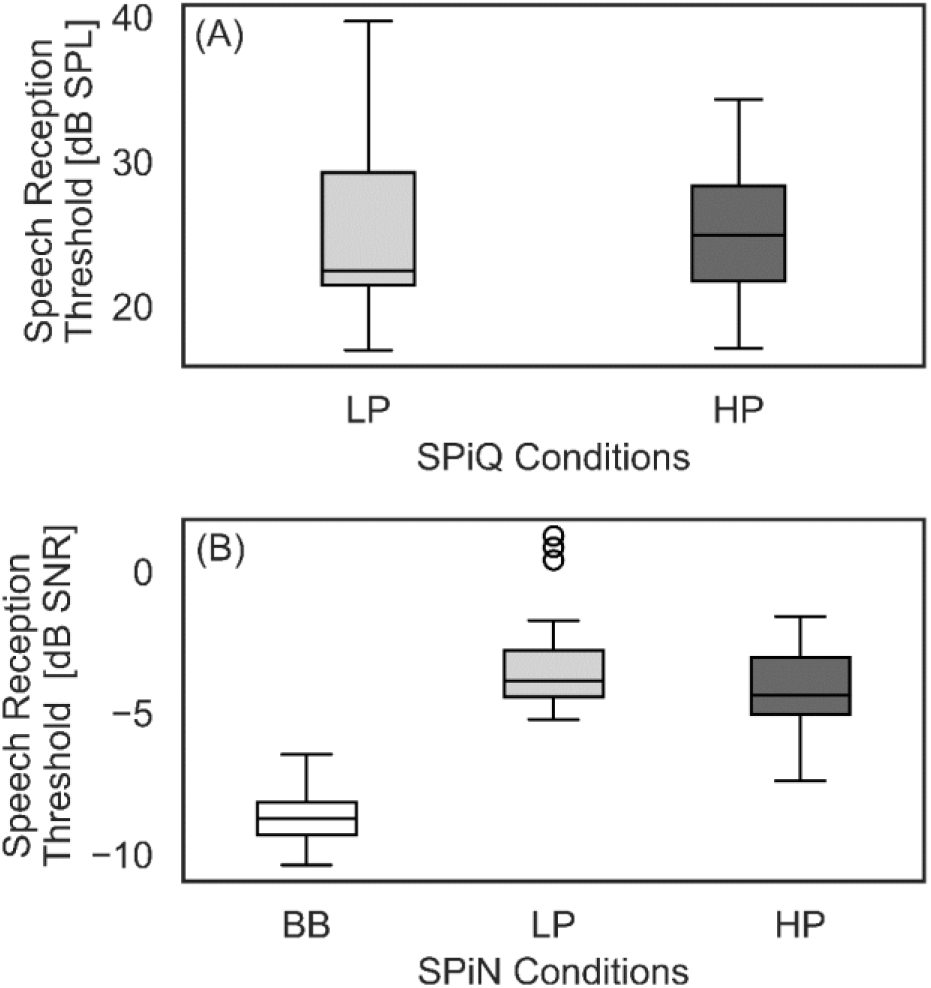
Boxplot of SPiQ (A) and SPiN (B) results at baseline session.

### BASELINE AEP-MEASUREMENTS

ABR latencies and amplitudes of waves I, III and V in the baseline session are given in Table 1. As expected from literature, latencies of all waves increased with lowering the stimulus level (i.e. 100dB peSPL vs 80dB peSPL), faster stimulus presentation (i.e. 120 Hz vs 11 Hz) or narrower stimulus-spectrum (i.e. TB vs click stimuli; Picton (2011)). A statistically significant relation with Laeq,life was found for the ABR wave V latency using a 11Hz 100dB click stimulus (ρ=0.66, p = 0.002). No other significant relations of ABR latency with Laeq,life were found. Spearman’s correlation coefficients ranged from 0.02 (p = 0.92) to 0.20 (p = 0.40) for wave I, from 0.01 (p = 0.99) to 0.37 (p = 0.11) for wave III and from 0.02 (p = 0.93) to 0.37 (p = 0.11) for wave V latencies.

**Table 1.** ABR data for ABR- and EFR responses of all stimulus types. Mean, SD, median and IQR are presented for ABR latencies and amplitudes of waves I, III and V and I/V amplitude ratio and EFR-strength.

| Stimulus |  |  | 11Hz<br>100dB<sub>peSPL</sub><br>click | 11Hz<br>80dB<sub>peSPL</sub><br>click | 120Hz<br>100dB<sub>peSPL</sub><br>click | 120Hz<br>80dB<sub>peSPL</sub><br>click | TB<sub>4kHz</sub> | TB<sub>1 kHz</sub> |
| --- | --- | --- | --- | --- | --- | --- | --- | --- |
| Latencies | I | mean | 2.67 | 3.32 | 3.14 | 3.70 | 3.96 | 4.36 |
|  |  | SD | 0.19 | 0.27 | 0.39 | 0.27 | 0.44 | 0.53 |
|  |  | median | 2.70 | 3.35 | 3.10 | 3.70 | 3.90 | 4.45 |
|  |  | IQR | 0.20 | 0.27 | 0.40 | 0.20 | 0.75 | 0.67 |
|  | III | mean | 4.94 | 5.46 | 5.47 | 6.01 | 6.15 | 6.48 |
|  |  | SD | 0.23 | 0.31 | 0.33 | 0.33 | 0.38 | 0.70 |
|  |  | median | 4.90 | 5.50 | 5.45 | 6.00 | 6.05 | 6.50 |
|  |  | IQR | 0.20 | 0.35 | 0.40 | 0.27 | 0.65 | 0.75 |
|  | V | mean | 6.73 | 7.18 | 7.32 | 7.90 | 7.94 | 5.57 |
|  |  | SD | 0.38 | 0.43 | 0.38 | 0.43 | 0.35 | 0.43 |
|  |  | median | 6.70 | 7.25 | 7.30 | 8.00 | 7.90 | 8.60 |
|  |  | IQR | 0.47 | 0.35 | 0.47 | 0.37 | 0.40 | 0.45 |
| Amplitudes | I | mean | 0.15 | 0.08 | 0.09 | 0.05 | 0.02 | 0.02 |
|  |  | SD | 0.10 | 0.06 | 0.06 | 0.04 | 0.04 | 0.05 |
|  |  | median | 0.17 | 0.07 | 0.08 | 0.04 | 0.03 | 0.02 |
|  |  | IQR | 0.14 | 0.11 | 0.08 | 0.09 | 0.05 | 0.06 |
|  | III | mean | 0.07 | 0.03 | 0.13 | 0.15 | 0.02 | -0.01 |
|  |  | SD | 0.07 | 0.07 | 0.10 | 0.09 | 0.06 | 0.07 |
|  |  | median | 0.05 | 0.03 | 0.11 | 0.14 | 0.01 | -0.01 |
|  |  | IQR | 0.13 | 0.10 | 0.13 | 0.15 | 0.05 | 0.10 |
|  | V | mean | 0.31 | 0.27 | 0.27 | 0.23 | 0.13 | 0.10 |
|  |  | SD | 0.10 | 0.11 | 0.11 | 0.08 | 0.08 | 0.07 |
|  |  | median | 0.30 | 0.27 | 0.26 | 0.22 | 0.13 | 0.09 |
|  |  | IQR | 0.12 | 0.10 | 0.13 | 0.08 | 0.09 | 0.10 |
| Amplitude ratios | I/V | mean | 0.47 | 0.34 | 0.39 | 0.22 | -0.96 | 0.44 |
|  |  | SD | 0.30 | 0.31 | 0.34 | 0.37 | 3.80 | 1.14 |
|  |  | median | 0.44 | 0.30 | 0.31 | 0.21 | 0.15 | 0.17 |
|  |  | IQR | 0.46 | 0.38 | 0.26 | 0.42 | 0.37 | 0.55 |
| Stimulus |  |  | SAM |  |  | RAM |  |  |
| EFR Magnitude (μV) |  | mean | 0.03 |  |  | 0.08 |  |  |
|  |  | SD | 0.01 |  |  | 0.03 |  |  |
|  |  | median | 0.03 |  |  | 0.09 |  |  |
|  |  | IQR | 0.01 |  |  | 0.04 |  |  |

Second, wave I, III and V ABR amplitudes showed no statistically significant monotonic relation to Laeq,life. For wave I, Spearman’s correlation coefficients of the different stimulus types ranged from −0.16 (p = 0.51) to 0.14 (p = 0.56), for wave III from −0.34 (p = 0.15) to 0.27 (p = 0.25) and for wave V from −0.25 (p = 0.29) to 0.003 (p = 0.99).

Finally, no significant correlation was found between the ABR I/V amplitude ratio and Laeq,life in all conditions. Spearman’s correlation coefficients ranged from −0.41 (p = 0.87) to 0.32 (p = 0.18) for the various ABR stimuli.

EFR magnitude of RAM and SAM stimuli in relation to Laeq,life is shown in Figure 6 (RAM: green circles SAM: gray circles). For all but one subject, higher EFR magnitudes were found using the RAM (mean 0.085µV, SD 0.029 µV, median 0.088 µV, range 0.007 – 0.136 µV) compared to the SAM-stimulus (mean 0.030 µV, SD 0.010 µV, median 0.028 µV, range 0.012 – 0.056 µV), consistent with auditory model predictions (Vasilkov et al., 2021). A negative, but statistically non-significant Spearman’s correlation coefficient was found in relation to Laeq,life for the SAM-stimulus (ρ=-0.34, p = 0.15) as well as for RAM-stimulus (ρ=-0.17, p = 0.49). When the two extreme cases— one with the lowest Laeq,life value (i.e. about 10 dB less than the other subjects) and another with the highest Laeq,life value (i.e. 10 dB higher than the rest of the cohort)— were omitted, respective Spearman’s correlation coefficients weakened (RAM-EFR: *ρ*=-0.12, p = 0.63, SAM-EFR: *ρ*=-0.09, p = 0.72).

**Figure 6:**
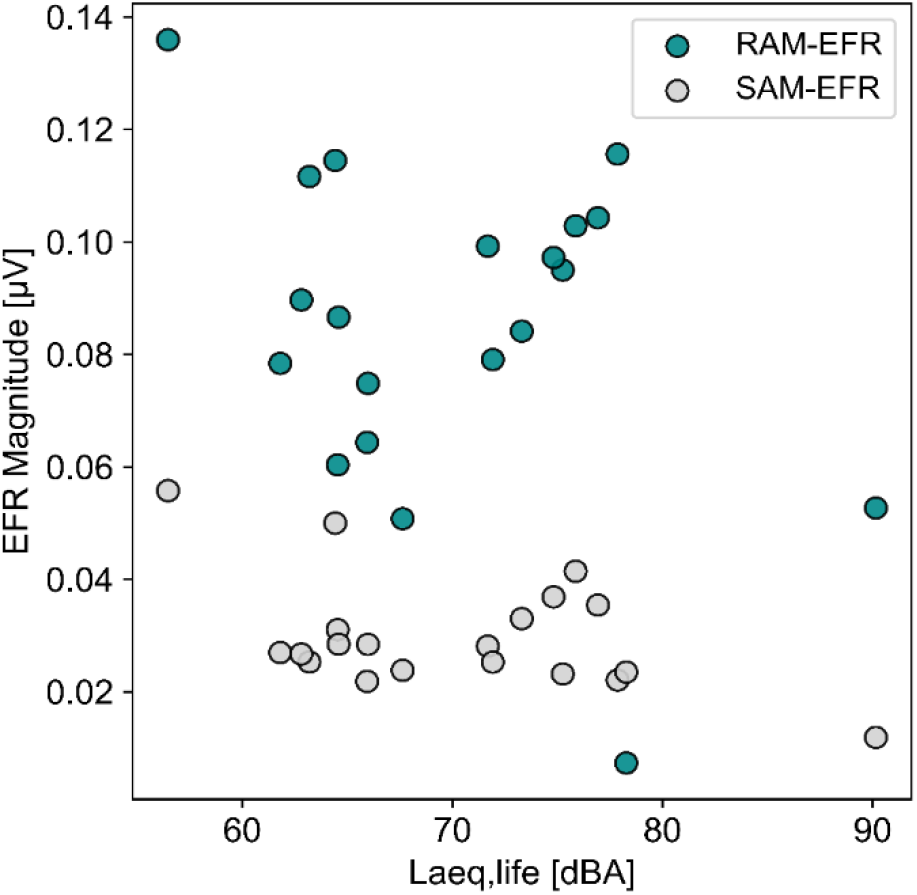
Scatterplot of EFR strength in relation to Laeq,life for SAM-stimuli (black triangles) and RAM-stimuli (grey triangles).

## RESULTS PART II

### QUESTIONNAIRE

The mean duration of the music festival attendance was 2.68 days (SD 1.29, range 1 - 5), where subjects self-reported to be exposed to music 8.9 hours a day (SD = 2.39, range 5 - 12). Estimation of noise-exposure levels during the festival, HPD-attenuation taken into account, resulted in an average Laeq, festival of 76.2 dBA (SD 7.82, range 59.35 – 88.19). Thirteen subjects (68.4%) reported to have worn HPDs during the music festivals and only 7 participants (36.8%) had worn them 50% of the time during noise exposure. Eight subjects (42.1%) experienced HRS after the noise exposure: six subjects (31.6%) experienced dullness after the music event, 2 persons (10.5%) experienced a subjective hearing loss and 3 persons (15.8%) reported tinnitus. Dullness and tinnitus were experienced both unilaterally or bilaterally and disappeared after a maximum of 24 hours after the noise-exposure, whereas a subjective hearing loss was experienced in both ears and disappeared after 30 minutes. Only one of the test subjects that had worn HPDs more than 50% of the noise-exposure time reported HRS. Baseline testing results of PTA, SPiQ and SPiN, and EFR did not differ significantly between subjects who reported HRS after the event, and subjects who did not. ABR results only showed between-group differences in the rate of 120 Hz, 100 dB click condition only. For this condition, the group with HRS had a significantly (p = 0.01) smaller wave-III latency (m 5.30 ms, range 4.60 – 5.50) than the group without HRS (m 5.70 ms, range 5.20 – 6.10). Second, wave-I amplitudes were significantly (p = 0.005) higher in the group with HRS (m 0.13 µV, range 0.02 – 0.22) than the group without (m 0.06 µV, range 0.03 – 0.09). Finally, a significantly higher I/V amplitude ratio was found in the HRS group (m 0.40, range 0.27 – 1.55) compared to the non-HRS group (m 0.19, range 0.09 – 0.49).

### LONGITUDINAL ANALYSIS OF THE HEARING STATUS

At single conventional audiogram frequency, a threshold elevation of 10 dB or more was found in eight subjects (42.1%) in the second session. At extended high frequencies, threshold shifts of 10 dB or higher were found in nine subjects (47.3%) between session 1 and session 2. All frequencies included, a threshold-shift of 10 dB or more was found in 15 subjects (78.9%). When defining a TTS as described by Occupational Safety and Health Administration (OSHA, 1974), 18 out of 19 subjects did not meet the TTS criterion. 18 subjects had an average threshold-shift from session 1 to session 2 at 2, 3 and 4 kHz, between −3.3 to 5 dB and did therefore not meet the average 10 dB TTS criterion at frequencies of 2, 3 and 4 kHz. However, in the second session, i.e. one day after the noise-exposure, an OSHA-defined TTS was found in one subject (0.05%). For this subject, an average elevation of 11.7 dB was found in session 2 compared to session 1, which recovered to values below 10 dB in sessions 3 and 4. A residual shift of 6.7 dB from session 1 was present in the fourth session. This subject experienced bilateral tinnitus (24h), but reported neither subjective hearing loss nor dullness.

Pure tone audiometry and χ^2^ results are displayed in Table 2. Significant intersession differences were found for auditory thresholds of 500 Hz (p = 0.033) and 10 kHz (p = 0.005). However, post-hoc analysis with Bonferroni adjustment showed no significant threshold-shift from the first session to a follow-up session (p>0.017). PTAs revealed statistically insignificant between-session differences (p>0.05). Even though subjects who experienced HRS showed a median positive shift, i.e. higher values, in PTA3-8kHz (3.13 dB HL, range −2.50 – 10.00) and PTA10-16kHz (3. 75 dB HL, range −3.75 – 6.25) in the second session and subjects without HRS did not (respectively 0.00 dB HL, range −3.75 – 3.75 and 0.00 dB HL, range −6.25 – 11.2). However, the between-group differences were not significant (p = 0.18 and 0.44, respectively).

**Table 2:** χ^2^-values of related-samples Friedman’s ANOVA for PTA. Significant results are highlighted with *(p<0.05), **(p<0.01) or ***(p<0.001). Median test values and IQRs are shown for each test session.

| Table 2: $\chi^2$-values of related-samples Friedman's ANOVA for PTA. Significant results are highlighted with *($p<0.05$), **($p<0.01$) or ***($p<0.001$). Median test values and IQRs are shown for each test session. | | | | | | | | | | |
| --- | --- | --- | --- | --- | --- | --- | --- | --- | --- | --- |
| Auditory thresholds (dB<sub>HL</sub>) | Frequency (Hz) | $\chi^2(3)$ | Session1 (pre) | | Session2 (1<sup>st</sup> post) | | Session3 (2<sup>nd</sup> post) | | Session4 (3<sup>d</sup> post) | |
|  |  |  | IQR | med | IQR | med | med | IQR | med | IQR |
|  | 125 | 5.56 | 5 | 5 | 10 | 10 | 5 | 10 | 5 | 10 |
|  | 250 | 3.47 | 5 | 10 | 5 | 5 | 5 | 10 | 5 | 10 |
|  | 500 | 8.73* | 0 | 5 | 5 | 10 | 0 | 5 | 0 | 5 |
|  | 1000 | 4.73 | 5 | 5 | 0 | 5 | 0 | 5 | 0 | 10 |
|  | 2000 | 0.26 | 0 | 10 | 0 | 10 | 0 | 10 | 0 | 10 |
|  | 3000 | 0.22 | 0 | 5 | 0 | 5 | 0 | 5 | 0 | 10 |
|  | 4000 | 2.83 | 0 | 5 | 0 | 5 | 0 | 5 | 0 | 10 |
|  | 6000 | 2.03 | 10 | 5 | 10 | 15 | 10 | 15 | 10 | 10 |
|  | 8000 | 6.33 | 10 | 5 | 10 | 10 | 5 | 10 | 10 | 10 |
|  | 10000 | 12.69** | 5 | 5 | 5 | 10 | 5 | 10 | 5 | 10 |
|  | 12000 | 4.88 | 5 | 15 | 10 | 20 | 5 | 15 | 5 | 15 |
|  | 14000 | 4.76 | 5 | 15 | 5 | 10 | 0 | 15 | 0 | 5 |
|  | 16000 | 7.39 | 10 | 30 | 10 | 20 | 10 | 25 | 5 | 25 |
|  | PTA<sub>3-8kHz</sub> | 3.84 | 5 | 5 | 5 | 8.75 | 3.75 | 5 | 5 | 8.75 |
|  | PTA<sub>10-16kHz</sub> | 6.97 | 2.5 | 12.5 | 7.5 | 13.75 | 6.25 | 10 | 5 | 7.5 |

### LONGITUDINAL ANALYSIS SPEECH AUDIOMETRY

Figure 7A displays the results of the SPiQ test. SPiQ showed slightly increased SRT-values from session 1 to session 2 for both LP- and HP-filtered speech conditions. Friedman’s ANOVA showed statistically significant intersession differences only for the HP filtered condition (χ^2^(3)=29.46, p<0.001). Although post-hoc analysis with Bonferroni-correction showed statistically insignificant changes between session 1 and session 2, a significant improvement (p = 0.008) in speech intelligibility from session 1 (med =22.67, IQR 8.04) to session 4 (med=20.78, IQR 5.28) was found.

**Figure 7:**
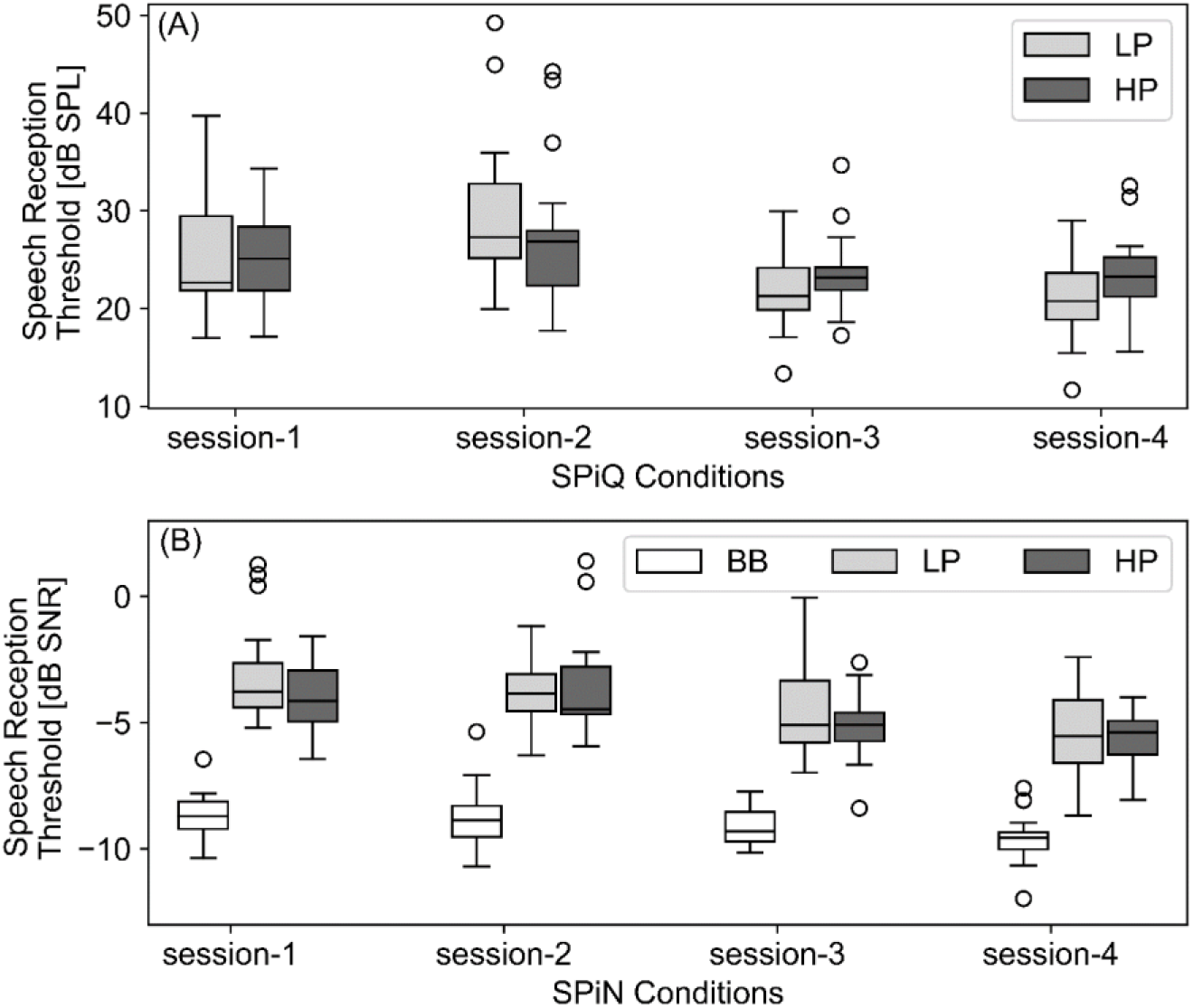
Boxplots of speech audiometry in quiet (A) and in noise (B) for all conditions and all sessions. Dark grey boxplots represent SRTs for BB stimuli, light grey and white boxplots represent SRTs for HP and LP conditions, respectively. Intersession differences, compared to session 1, were significant if p<0.017 and are highlighted with one, two or three asterisks (resp. p<0.017, p<0.003 and p<0.0003). Both SPiN and SPIQ showed no significant differences between sessions 1 and 2.

The same trend of improved speech intelligibility over sessions was found for SPiN, as shown in Figure 7B. Friedman’s ANOVA showed statistically significant intersession differences for the BB as well as the LP condition (χ^2^(3)=29.72, p<0.001) and the HP condition (χ^2^(3)=27.97, p<0.001). Post-hoc pairwise comparisons with Bonferroni-correction showed a significant improvement of SNR from session 1 to 4 for all conditions (BB p = 0.002, LP and HP p<0.001) and a significant improvement from session 1 to 3 for the LP condition (p = 0.006). However, no statistical significant differences between session 1 and session 2 were found for all test conditions.

Between-group comparisons of persons with or without HRS, did not show significant differences in SRT-shift from session1 to session 2 for both SPiQ (p-range of 0.11 to 0.41) and SPiN (p-range of 0.06 to 0.77).

### LONGITUDINAL ANALYSIS AEP-MEASUREMENTS

Table 3 represents all results and χ^2^-values of AEP-measurements. ABR-latencies showed statistically insignificant changes between sessions for any wave or stimulus type. ABR wave III showed significant intersession differences only for the 11-Hz 100-dB peSPL stimulus (χ^2^(3)=7.93, p = 0.48) and wave V changed significantly over time for the 11-Hz 80-dB peSPL stimulus (χ^2^(3)=12.92, p = 0.005), but post-hoc comparisons with Bonferroni correction showed no significant differences between the first session and sessions 2, 3 or 4 (p>0.017). Amplitude ratios I/V did not reveal any significant intersession changes for the considered stimulus types (p>0.05).

**Table 3:**
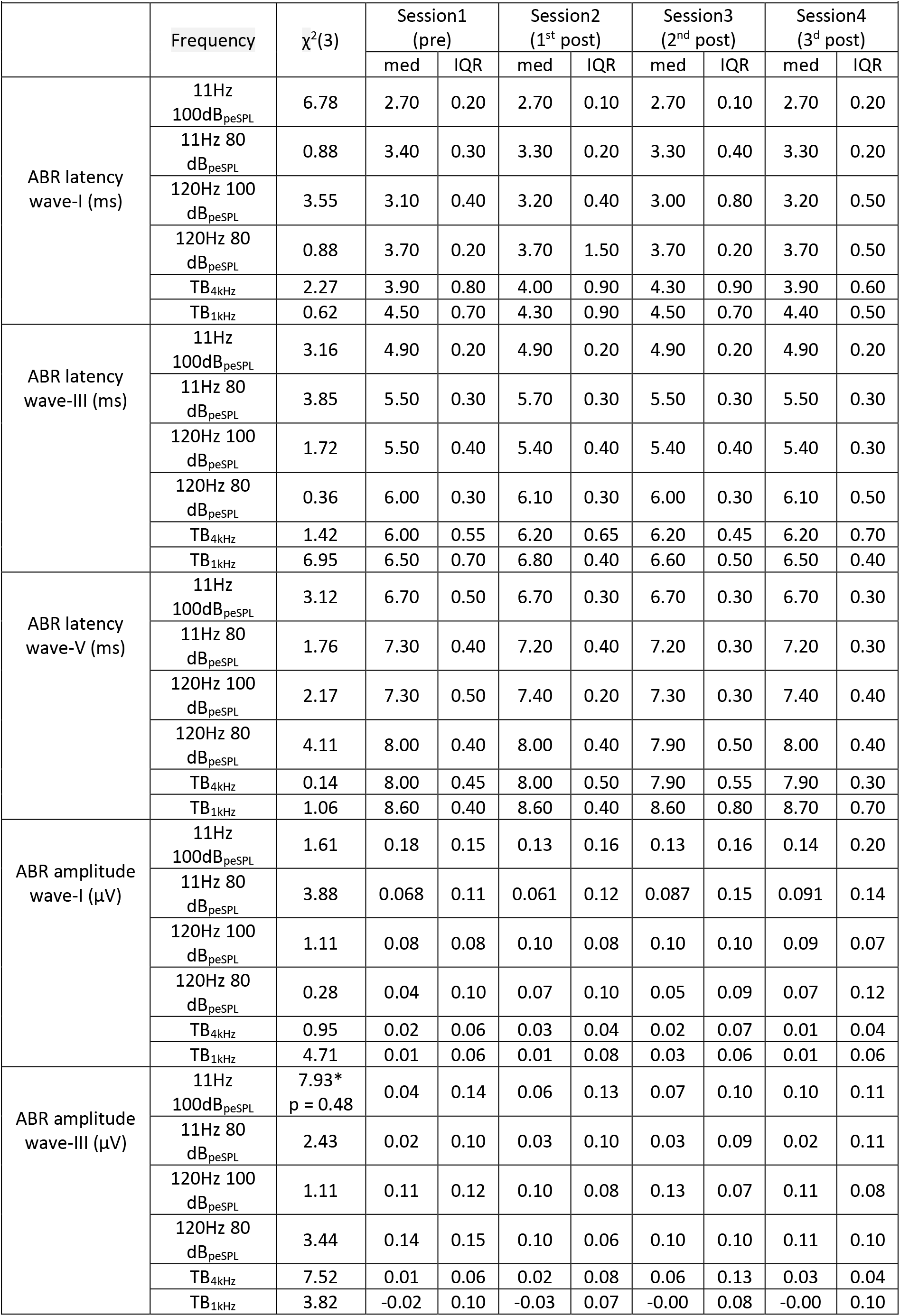
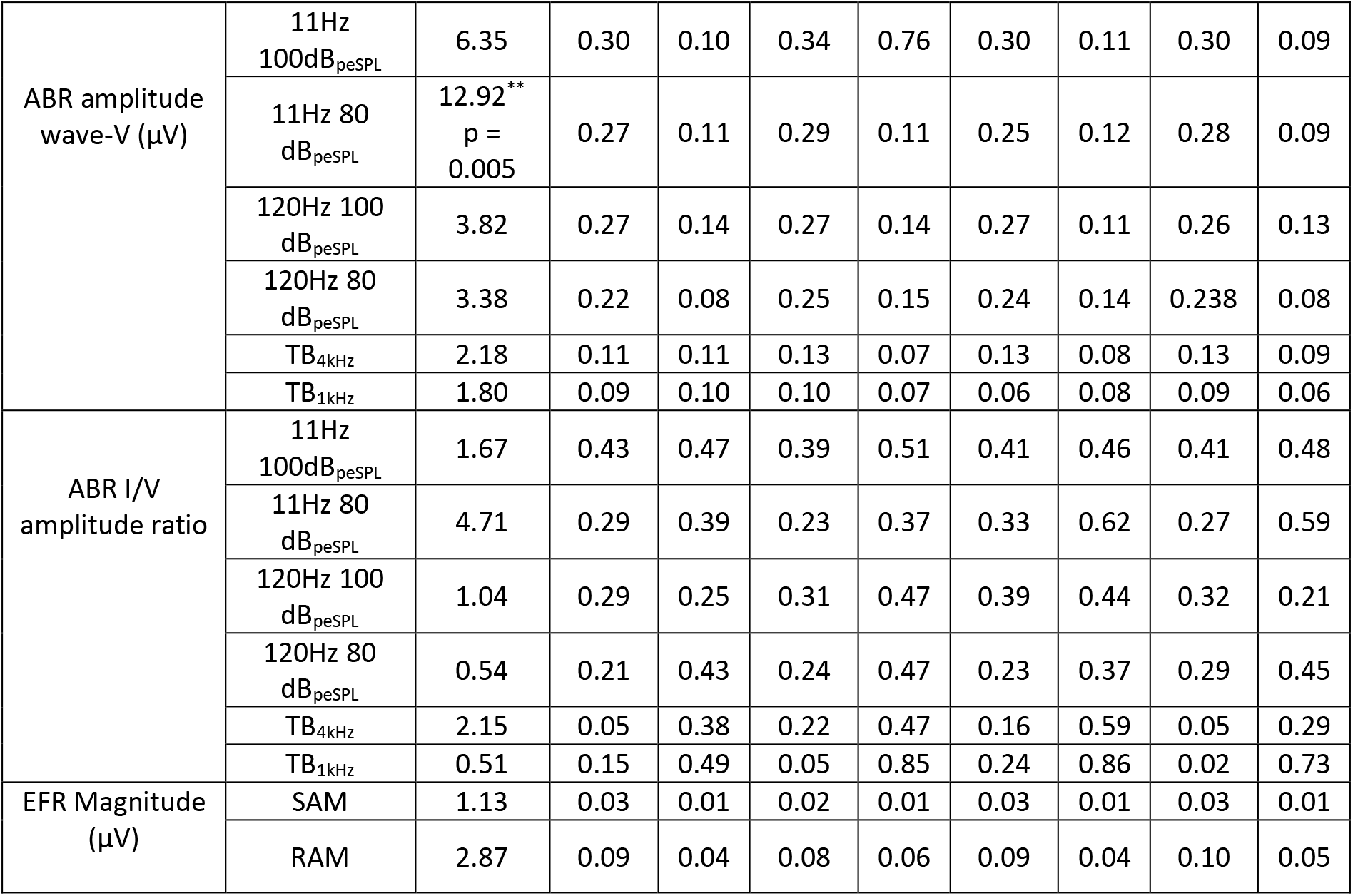
χ^2^-values of related-samples Friedman’s ANOVA for AEP measurements. Significant results are highlighted with *(p<0.05), **(p<0.01) or ***(p<0.001). Median test values and IQRs are shown for each test session.

| Table 3: $\chi^2$-values of related-samples Friedman's ANOVA for AEP measurements. Significant results are highlighted with *($p<0.05$), **($p<0.01$) or ***($p<0.001$). Median test values and IQRs are shown for each test session. | | | | | | | | | | |
| --- | --- | --- | --- | --- | --- | --- | --- | --- | --- | --- |
| | Frequency | $\chi^2(3)$ | Session1<br>(pre) | | Session2<br>(1<sup>st</sup> post) | | Session3<br>(2<sup>nd</sup> post) | | Session4<br>(3<sup>d</sup> post) | |
|  |  |  | med | IQR | med | IQR | med | IQR | med | IQR |
| ABR latency<br>wave-I (ms) | 11Hz<br>100dB<sub>peSPL</sub> | 6.78 | 2.70 | 0.20 | 2.70 | 0.10 | 2.70 | 0.10 | 2.70 | 0.20 |
|  | 11Hz 80<br>dB<sub>peSPL</sub> | 0.88 | 3.40 | 0.30 | 3.30 | 0.20 | 3.30 | 0.40 | 3.30 | 0.20 |
|  | 120Hz 100<br>dB<sub>peSPL</sub> | 3.55 | 3.10 | 0.40 | 3.20 | 0.40 | 3.00 | 0.80 | 3.20 | 0.50 |
|  | 120Hz 80<br>dB<sub>peSPL</sub> | 0.88 | 3.70 | 0.20 | 3.70 | 1.50 | 3.70 | 0.20 | 3.70 | 0.50 |
|  | TB<sub>4kHz</sub> | 2.27 | 3.90 | 0.80 | 4.00 | 0.90 | 4.30 | 0.90 | 3.90 | 0.60 |
|  | TB<sub>1kHz</sub> | 0.62 | 4.50 | 0.70 | 4.30 | 0.90 | 4.50 | 0.70 | 4.40 | 0.50 |
| ABR latency<br>wave-III (ms) | 11Hz<br>100dB<sub>peSPL</sub> | 3.16 | 4.90 | 0.20 | 4.90 | 0.20 | 4.90 | 0.20 | 4.90 | 0.20 |
|  | 11Hz 80<br>dB<sub>peSPL</sub> | 3.85 | 5.50 | 0.30 | 5.70 | 0.30 | 5.50 | 0.30 | 5.50 | 0.30 |
|  | 120Hz 100<br>dB<sub>peSPL</sub> | 1.72 | 5.50 | 0.40 | 5.40 | 0.40 | 5.40 | 0.40 | 5.40 | 0.30 |
|  | 120Hz 80<br>dB<sub>peSPL</sub> | 0.36 | 6.00 | 0.30 | 6.10 | 0.30 | 6.00 | 0.30 | 6.10 | 0.50 |
|  | TB<sub>4kHz</sub> | 1.42 | 6.00 | 0.55 | 6.20 | 0.65 | 6.20 | 0.45 | 6.20 | 0.70 |
|  | TB<sub>1kHz</sub> | 6.95 | 6.50 | 0.70 | 6.80 | 0.40 | 6.60 | 0.50 | 6.50 | 0.40 |
| ABR latency<br>wave-V (ms) | 11Hz<br>100dB<sub>peSPL</sub> | 3.12 | 6.70 | 0.50 | 6.70 | 0.30 | 6.70 | 0.30 | 6.70 | 0.30 |
|  | 11Hz 80<br>dB<sub>peSPL</sub> | 1.76 | 7.30 | 0.40 | 7.20 | 0.40 | 7.20 | 0.30 | 7.20 | 0.30 |
|  | 120Hz 100<br>dB<sub>peSPL</sub> | 2.17 | 7.30 | 0.50 | 7.40 | 0.20 | 7.30 | 0.30 | 7.40 | 0.40 |
|  | 120Hz 80<br>dB<sub>peSPL</sub> | 4.11 | 8.00 | 0.40 | 8.00 | 0.40 | 7.90 | 0.50 | 8.00 | 0.40 |
|  | TB<sub>4kHz</sub> | 0.14 | 8.00 | 0.45 | 8.00 | 0.50 | 7.90 | 0.55 | 7.90 | 0.30 |
|  | TB<sub>1kHz</sub> | 1.06 | 8.60 | 0.40 | 8.60 | 0.40 | 8.60 | 0.80 | 8.70 | 0.70 |
| ABR amplitude<br>wave-I (μV) | 11Hz<br>100dB<sub>peSPL</sub> | 1.61 | 0.18 | 0.15 | 0.13 | 0.16 | 0.13 | 0.16 | 0.14 | 0.20 |
|  | 11Hz 80<br>dB<sub>peSPL</sub> | 3.88 | 0.068 | 0.11 | 0.061 | 0.12 | 0.087 | 0.15 | 0.091 | 0.14 |
|  | 120Hz 100<br>dB<sub>peSPL</sub> | 1.11 | 0.08 | 0.08 | 0.10 | 0.08 | 0.10 | 0.10 | 0.09 | 0.07 |
|  | 120Hz 80<br>dB<sub>peSPL</sub> | 0.28 | 0.04 | 0.10 | 0.07 | 0.10 | 0.05 | 0.09 | 0.07 | 0.12 |
|  | TB<sub>4kHz</sub> | 0.95 | 0.02 | 0.06 | 0.03 | 0.04 | 0.02 | 0.07 | 0.01 | 0.04 |
|  | TB<sub>1kHz</sub> | 4.71 | 0.01 | 0.06 | 0.01 | 0.08 | 0.03 | 0.06 | 0.01 | 0.06 |
| ABR amplitude<br>wave-III (μV) | 11Hz<br>100dB<sub>peSPL</sub> | 7.93*<br>p = 0.48 | 0.04 | 0.14 | 0.06 | 0.13 | 0.07 | 0.10 | 0.10 | 0.11 |
|  | 11Hz 80<br>dB<sub>peSPL</sub> | 2.43 | 0.02 | 0.10 | 0.03 | 0.10 | 0.03 | 0.09 | 0.02 | 0.11 |
|  | 120Hz 100<br>dB<sub>peSPL</sub> | 1.11 | 0.11 | 0.12 | 0.10 | 0.08 | 0.13 | 0.07 | 0.11 | 0.08 |
|  | 120Hz 80<br>dB<sub>peSPL</sub> | 3.44 | 0.14 | 0.15 | 0.10 | 0.06 | 0.10 | 0.10 | 0.11 | 0.10 |
|  | TB<sub>4kHz</sub> | 7.52 | 0.01 | 0.06 | 0.02 | 0.08 | 0.06 | 0.13 | 0.03 | 0.04 |
|  | TB<sub>1kHz</sub> | 3.82 | -0.02 | 0.10 | -0.03 | 0.07 | -0.00 | 0.08 | -0.00 | 0.10 |
| ABR amplitude wave-V ($\mu\text{V}$) | 11Hz 100dB<sub>peSPL</sub> | 6.35 | 0.30 | 0.10 | 0.34 | 0.76 | 0.30 | 0.11 | 0.30 | 0.09 |
|  | 11Hz 80 dB<sub>peSPL</sub> | 12.92**<br>p = 0.005 | 0.27 | 0.11 | 0.29 | 0.11 | 0.25 | 0.12 | 0.28 | 0.09 |
|  | 120Hz 100 dB<sub>peSPL</sub> | 3.82 | 0.27 | 0.14 | 0.27 | 0.14 | 0.27 | 0.11 | 0.26 | 0.13 |
|  | 120Hz 80 dB<sub>peSPL</sub> | 3.38 | 0.22 | 0.08 | 0.25 | 0.15 | 0.24 | 0.14 | 0.238 | 0.08 |
|  | TB<sub>4kHz</sub> | 2.18 | 0.11 | 0.11 | 0.13 | 0.07 | 0.13 | 0.08 | 0.13 | 0.09 |
|  | TB<sub>1kHz</sub> | 1.80 | 0.09 | 0.10 | 0.10 | 0.07 | 0.06 | 0.08 | 0.09 | 0.06 |
| ABR I/V amplitude ratio | 11Hz 100dB<sub>peSPL</sub> | 1.67 | 0.43 | 0.47 | 0.39 | 0.51 | 0.41 | 0.46 | 0.41 | 0.48 |
|  | 11Hz 80 dB<sub>peSPL</sub> | 4.71 | 0.29 | 0.39 | 0.23 | 0.37 | 0.33 | 0.62 | 0.27 | 0.59 |
|  | 120Hz 100 dB<sub>peSPL</sub> | 1.04 | 0.29 | 0.25 | 0.31 | 0.47 | 0.39 | 0.44 | 0.32 | 0.21 |
|  | 120Hz 80 dB<sub>peSPL</sub> | 0.54 | 0.21 | 0.43 | 0.24 | 0.47 | 0.23 | 0.37 | 0.29 | 0.45 |
|  | TB<sub>4kHz</sub> | 2.15 | 0.05 | 0.38 | 0.22 | 0.47 | 0.16 | 0.59 | 0.05 | 0.29 |
|  | TB<sub>1kHz</sub> | 0.51 | 0.15 | 0.49 | 0.05 | 0.85 | 0.24 | 0.86 | 0.02 | 0.73 |
| EFR Magnitude ($\mu\text{V}$) | SAM | 1.13 | 0.03 | 0.01 | 0.02 | 0.01 | 0.03 | 0.01 | 0.03 | 0.01 |
|  | RAM | 2.87 | 0.09 | 0.04 | 0.08 | 0.06 | 0.09 | 0.04 | 0.10 | 0.05 |

RAM-EFR magnitudes were consistently larger than SAM-EFR magnitudes. Even though the median EFR magnitude showed a slight reduction from the first to the second session for both SAM and RAM stimuli, the EFR magnitude did not significantly change across sessions (p>0.05).

ABR-shifts from session 1 to session 2, compared between groups who did or did not experience HRS, respectively, can be found in Table 1 of the Supplemental Material. Shifts in ABR latencies, amplitudes and I/V amplitude showed some diffuse significant group differences for certain stimuli, e.g. ABR wave I latency shift for 11-Hz 80-dB click stimulus (p = 0.02), wave-III latency (p = 0.04), ABR wave-I (p = 0.04) and wave-V (p = 0.02) amplitude shift for TB 4-kHz stimulus, and I/V-amplitude ratio for the 120-Hz 100-dB stimulus (p = 0.03). However, no consistent trends across different stimulus types were observed.

The EFR magnitude did show a negative shift, i.e. an EFR-strength reduction from session 1 to session 2 for both stimulus types within the group that reported HRS (SAM: med = −0.004 range = −0.03 - 0.02; RAM: med = −0.007; range = −0.06 – 0.05). In the group without HRS, a negative shift was only found with the SAM stimulus (med = −0.006; range = −0.01 – 0.1), but not for the RAM stimulus (med = 0.008; range = −0.06 – 0.03). No significant group differences were found between groups with or without HRS in EFR magnitude shifts, neither for SAM (U = 49.00, p = 0.71) nor for RAM-stimuli (U = 52.00, p = 0.88). Figure 5 in Supplemental Material shows the Spearman’s correlation between Laeq,festival and RAM and SAM-EFR magnitudes, ABR (80 dB-peSPL, 11 Hz) wave I and V amplitudes and latencies and PTA10-16kHz (averaged EHF audiogram).

## DISCUSSION

### PART I

The first aim of this study consisted of a baseline evaluation of hearing status in relation to the noise-exposure history of 19 young, normal-hearing adults. Our inclusion criteria, specifically normal auditory thresholds and the absence of HRS, eliminated ears with OHC-dysfunction as much as possible.

As other studies failed to find a relation between young adults’ hearing thresholds or distortion product otoacoustic emissions (DPOAEs) and their lifetime noise-exposure history (Degeest, Clays, et al., 2017; Keppler, Dhooge, et al., 2015b; Wei et al., 2017; Williams et al., 2015), no substantial OHC-loss due to noise-exposure was expected in this population. Indeed, PTA10-16kHz showed no significant correlation with lifetime noise-exposure history, indicating relatively intact functioning of the OHCs. Nevertheless, in future studies, DPOAE measurements can still provide a tool to precisely assess and monitor the integrity of OHCs in the apical cochlear regions.

The assumption of a normally functioning cochlear amplifier facilitates interpretation of the other hearing test outcomes in the context of hidden hearing loss or CS. CS has been linked to supra-threshold hearing function (Kujawa & Liberman, 2009; Lin et al., 2011) and is thought to affect speech understanding, especially in difficult listening situations. Therefore, it could be hypothesized that SPiQ- or SPiN-metrics would relate to lifetime noise-exposure history if CS was present. However, no relation was found in this study population. This supports the results of the majority of studies investigating speech intelligibility in relation to noise-exposure history of normal-hearing persons, where no relation was found either (Couth et al., 2020; Fulbright et al., 2017; Grinn et al., 2017; Guest et al., 2018; Prendergast, Millman, et al., 2017; Yeend et al., 2017). In contrast, some studies have reported a link between noise-exposure history and speech intelligibility (Hope et al., 2013; Kumar et al., 2012; Liberman et al., 2016). The inclusion of subjects that were regularly exposed to occupational noise or a music education could have contributed to a higher noise-exposure dose in these latter studies and increased the risk of CS in the tested population. However, OHC damage could not be ruled out in all these studies since, respectively, audiometric thresholds were only measured up to 4kHz (Hope et al., 2013), or speech intelligibility deterioration was accompanied with audiometric loss in EHFs (Liberman et al., 2016). A recent study by Prendergast et al. (2019) found a link between noise-exposure history and speech intelligibility albeit showing a better performance in subjects with higher lifetime noise-exposure history at levels above 85 dBA. As many of their highly-exposed subjects worked in the music industry, listening strategies and attention training on complex soundscapes are proposed by the authors to explain this difference.

The baseline AEP-measurements in the present study did not show significant correlations to lifetime noise-exposure history. First, ABR wave I, III and V amplitudes and the ABR I/V amplitude ratio were not significantly correlated to noise-exposure history. These observations corroborate results from a number of human studies that did not show a clear link between ABR wave I amplitudes and recreational noise-exposure (Couth et al., 2020; Fulbright et al., 2017; Grinn et al., 2017; Liberman et al., 2016; Prendergast et al., 2018; Ridley et al., 2018; Skoe & Tufts, 2018; Spankovich, Le Prell, et al., 2017). However, two studies performed by Bramhall et al. (2017) and Stamper and Johnson (2015) revealed significant wave I amplitude reductions in subjects with higher noise-exposure history, although for the latter study this was only true for female subjects (Stamper & Johnson, 2015). Furthermore, Valderrama et al. (2018) reported reduced wave I amplitudes in relation to recreational and occupational noise exposure, but this relation was not significant after removal of outliers. Amplitudes of wave III have less commonly been analyzed and have not shown significant relations to noise-exposure (Ridley et al., 2018; Skoe & Tufts, 2018). Furthermore, wave V amplitudes did not show a relation to recreational noise exposure in literature (Prendergast et al., 2018; Ridley et al., 2018; Skoe & Tufts, 2018; Spankovich, Bishop, et al., 2017) and neither did I/V amplitude ratios (Prendergast, Guest, et al., 2017; Spankovich, Bishop, et al., 2017). One study did report a reduction in I/V ratios in subjects that frequently attended music concerts (Grose et al., 2019). On the contrary, a recent study by Couth et al. (2020) showed higher I/V-amplitude ratios in musicians compared to non-musicians. Even though noise-exposure history was similar between both groups in that study and noise-exposure history did not show a significant relation to I/V amplitude, the highly-exposed subjects showed OHC-dysfunction.

Second, regarding ABR latencies, no relation to noise-exposure history was found in the present study. This is in line with several studies, where ABR wave I latencies (Couth et al., 2020; Spankovich, Bishop, et al., 2017) or wave V latencies (Spankovich, Bishop, et al., 2017) did not relate to recreational noise-exposure. On the contrary, Skoe and Tufts (2018) did find a delay in waves I, III and V for subjects with a higher noise-exposure based on one-week dosimetry measurements. The more highly-exposed subjects in this study were mostly students who participated in music ensembles on campus. A delay in wave V latency with increasing noise-exposure was also found by Couth et al. (2020), but only for males, and by Prendergast, Guest, et al. (2017) where significance disappeared after correcting for age.

Finally, EFR magnitude did not correlate to lifetime noise-exposure history in the present study. This is in line with other studies in normal-hearing subjects (Grose et al., 2017) and in normal-hearing subjects with tinnitus (Guest et al., 2017). Prendergast, Guest, et al. (2017) found a significant negative correlation between EFR magnitude and noise-exposure. However, this was only true for males and after controlling for age, this correlation was no longer significant. A study by the same lab could not find noise-exposure as a predictor of EFR magnitude (Prendergast et al., 2019). In contrast, animal studies (Möhrle et al., 2016; Parthasarathy and Kujawa, 2018; Shaheen et al., 2015) and those that used computational modelling of the auditory periphery or human data have shown the possible value of EFRs in diagnosing CS (Bharadwaj et al., 2015; Keshishzadeh et al., 2020; Keshishzadeh et al., 2021; Paul et al., 2017; Vasilkov et al., 2021).

As described above, studies trying to demonstrate the presence of CS in humans by relating possible biomarkers to noise-exposure history, e.g. SPiN, ABR-amplitudes or amplitude ratios, ABR-latencies or EFR magnitude, report contradictory results (see Bramhall et al., 2019 for review). The disagreement between those studies can be caused by multiple reasons. First, study population characteristics such as age can vary across studies. Age-related CS could be dominant over noise-induced CS and therefore hiding the effects of noise-exposure history. The use of different age ranges between studies implies that in some study cohorts mostly occupational noise-exposure was studied (Hope et al., 2013; Kumar et al., 2012; Prendergast et al., 2019; Valderrama et al., 2018), whereas other studies focused on recreational noise-exposure. Second, methodological differences could lead to different conclusions. Based on (the absence of) EHF audiometry- or OAE-results, OHC-deficits cannot be ruled out in some study cohorts (Liberman et al., 2016). Furthermore, methods to calculate or measure noise-exposure history differ between studies. As objective measurements of noise-exposure such as dosimetry is hard to be collected during a lifetime period, subjective and retrospective questionnaires or interviews are used to estimate noise-exposure history. Beach et al. (2012) stated that realistic loudness estimations can be made through/via questionnaires about recent noise-exposures when compared to dosimetry metrics. However, questionnaires mostly do not take into account frequency components of noise or acoustic traumas caused by impulse noises. Other factors such as blood type (Doğru et al., 2003), nutrition (Kopke et al., 2005), fitness (Kolkhorst et al., 1998), smoking (Ferrite & Santana, 2005; Pouryaghoub et al., 2007), alcohol use (Kraaijenga et al., 2018; Upile et al., 2007) and drug use (Church et al., 2013) have been suggested to influence susceptibility to NIHL, but have not been taken into account in studies.

### PART II

The second part of this study evaluated longitudinal biomarker changes before and after attending a music festival.

Eight subjects (42.1%) experienced HRS after the music festival, comparable to Mercier et al. (2003), where 36% experienced HRS after attending a music festival. It has to be noted that the majority of subjects wore HPDs, at least for some time during the concerts. However, of those subjects who reported using HPDs, only a few wore them consistently. Only one of the subjects who wore HPDs over 50% of the time during noise-exposure experienced HRS, which is in line with findings that HPDs reduce HRS (Kraaijenga et al., 2018) or reduce TTS (Ramakers et al., 2016). Baseline testing results of subjects that reported HRS after the event, did not significantly differ from those who did not, suggesting that cross-sectional use of the present test battery does not predict the risk of developing symptoms such as tinnitus after noise-exposure.

A TTS as defined by OSHA-standards (OSHA, 1974) was only found in one subject one day after the music festival. No significant group changes in auditory thresholds were found for any of the tested frequencies. The difficulty of detecting a TTS in non-laboratory situations such as attending concerts or nightclubs was earlier described (Le Prell et al., 2012). The experienced noise-exposure is known to vary considerably between subjects attending such events, as noise intensity levels vary depending on distance relative to the stage and the music genre (Opperman et al., 2006). Furthermore, it is difficult to distinguish a small TTS from test-retest variability (Le Prell et al., 2012; Schlauch & Carney, 2012), which is amongst others dependent on participant experience and motivation (Schlauch & Carney, 2012). Above studies found a TTS in the majority of subjects after attending concerts, festivals or music venues (Emmerich et al., 2002; Le Prell, 2019; Opperman et al., 2006; Ramakers et al., 2016). These studies evaluated hearing thresholds immediately after noise-exposure, whereas the greatest auditory-threshold recovery was found to occur within two-to-four hours after the noise-exposure (Emmerich et al., 2002; Grinn et al., 2017; Le Prell et al., 2012). Therefore the possibility of a recovered TTS in the present study population should be considered, especially in those subjects that experienced HRS (Masterson et al., 2016; Schmuzigert et al., 2006).

SPiQ and SPiN tests showed no significant differences one day after noise exposure compared to the baseline session. This finding is in contrast with the study of Grinn et al. (2017), where the noise dose of a noise-exposure event has been related to a decrease in speech in noise intelligibility. However, this study showed a significant improvement in SRT-values for session 4 compared to session 1 for each condition, which could be related to a learning effect. Two training lists were used as suggested in literature concerning the Flemish Matrix speech material to overcome the largest learning effect. However, smaller and non-significant improvement in SRT-values are described even after the presentation of two training lists (Kollmeier et al., 2015). As a consequence of this learning effect, possible deteriorations in speech intelligibility shortly after noise-exposure could remain undetected.

No significant changes could be found between the first session and sessions 2, 3 or 4 for AEP-measurements, as determined by analysis of ABR amplitudes and latencies of wave I, III and V and ABR I/V amplitude ratio and EFR magnitudes. Grinn et al. (2017), who measured ABR wave I amplitudes by conducting electrocochleography, did not find amplitude reductions after attending a noisy event.

### DETECTING AND MONITORING CS IN NORMAL HEARING SUBJECTS

In the present study, no relation was found between noise-exposure history and possible biomarkers of CS, nor did those biomarkers change across the sessions which surrounded a music event in normal-hearing young adults. Baseline testing results and shifts from session 1 to session 2 did not differ between subjects that experienced HRS after the music event, and those who did not. Thus, no strong conclusions can be made regarding the presence of noise-induced CS in relation to noise-exposure history nor regarding the appearance of CS shortly after visiting a music festival in this cohort of young normal-hearing subjects.

Possibly, the noise-exposure history and subjectively reported recreational noise-exposure dose in the present study (mean: 76.2 dBA) were not sufficiently high to cause CS, despite the (100dBA, 1-hour) and (102dBA, 15 minutes) emission possibility at the considered festivals (Tronstad & Gelderblom, 2016). Permanent consequences of recreational noise on hearing has been doubted before, as reviewed by Carter et al. (2014). This theory is reflected by the absence of TTS (OSHA defined) in all but one subjects at one day after the festival. In noise-exposed animals, smaller TTSs of 20 – 30 dB, measured at one day after a single noise-exposure, were shown not to result in CS (Fernandez et al., 2015; Jensen et al., 2015), whereas TTSs up to 50 dB were reported for noise-exposed animals with substantial CS (Furman et al., 2013; Kujawa & Liberman, 2009; Lin et al., 2011). It may hence be that the young normal-hearing listeners we considered did not suffer from CS. Second, it is possible that the biomarkers for CS are not sensitive or specific enough to detect or to monitor CS in normal-hearing young adults. The interpretation of speech-in-noise audiometry is complicated by a possible learning effect.

In order to distinguish between the absence of CS in the study cohort and a lack of specificity in the CS biomarkers, future studies may provide more insight. First, it would be interesting to further investigate inter-subject variability and test-retest reliability of CS biomarkers, especially the recently developed RAM-EFR metric, in a cohort of normal-hearing subjects. Second, the use of dosimetry, where noise doses can be measured for specific noise events, logging of HPD-use during a music event, and hearing process evaluation immediately after the event will provide the needed information regarding recent noise-exposure doses, HPD-use and temporary versus long-term effects on hearing. Given the difficulties associated with equipping the subjects with microphones or dosimeters to measure the festival noise-level, another possible option could be to perform noise measurements at the festival and take sample measures of the noise-level at different distances from the stage/loudspeakers. The subsequent analysis could then consider the average of those measures together with the duration of the subjects attendance to the event. Third, as AEP-measurements can be conducted with different stimulus parameters and rates, their sensitivity to detect CS can be further investigated by further comparing their results within the same study cohorts (Garrett et al., 2020; Vasilkov et al., 2021). Finally, larger test groups will allow us to compare different possibly confounding factors such as the use of HPD and different levels of noise-exposure history with more statistical power.

## ACKNOWLEDGMENT

This work was supported by UGent BOF-IOP project: “Portable Hearing Diagnostics: Monitoring of Auditory-nerve Integrity after Noise Exposure (EarDiMon)” (T.V.M., N.D.P., I.D., H.K. and S.V.) and European Research Council (ERC) under the Horizon 2020 Research and Innovation Program, grant agreement no. 678120 RobSpear (S.K. and S.V.).

## Supplemental Material

This section provides additional information on the relation between Laeq,life and EHF audiometric thresholds of different (Figure 1) and averaged frequencies (Figure 2), the variation of speech reception thresholds of different conditions in relation to Laeq,life (Figure 3 and 4) and correlation between Laeq,festival and shifted amount of different AEP markers as well as EHF audiogram from session 1 to session 2 (Figure 5). Furthermore, the amplitude and latency shifts of ABR wave I and V from session 1 to session 2 are reported in Table 1. The shifted amounts are specified separately for groups who did or did not experience HRS after attending the music-event.

**Figure 8.**
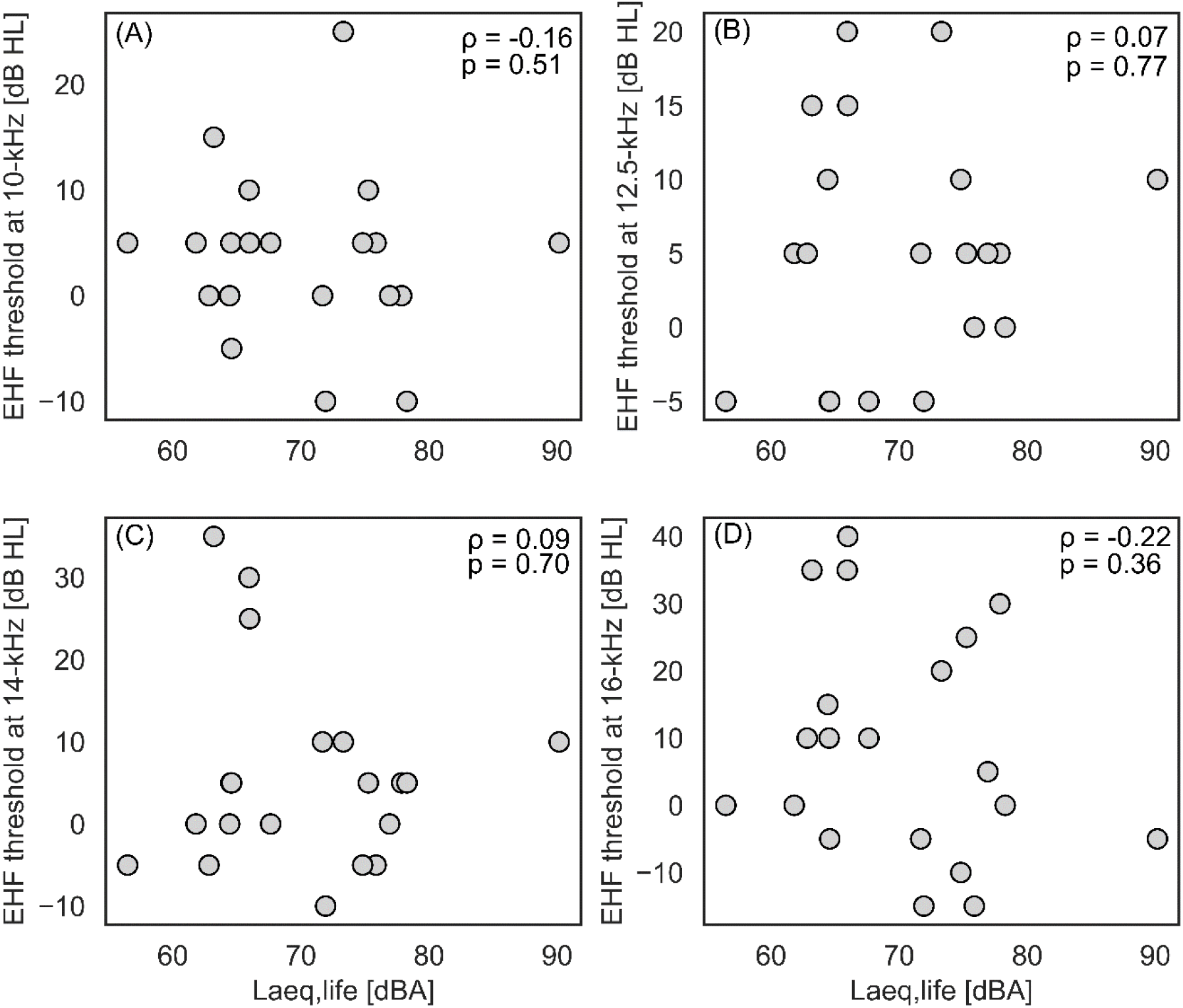
Spearman’s correlation analysis between Laeq,life and (A) 10 kHz, (B) 12.5 kHz, (C) 14 kHz and (D) 16 kHz EHF thresholds. Correlation coefficients and p-values are indicated in respective panels.

**Figure 9.**
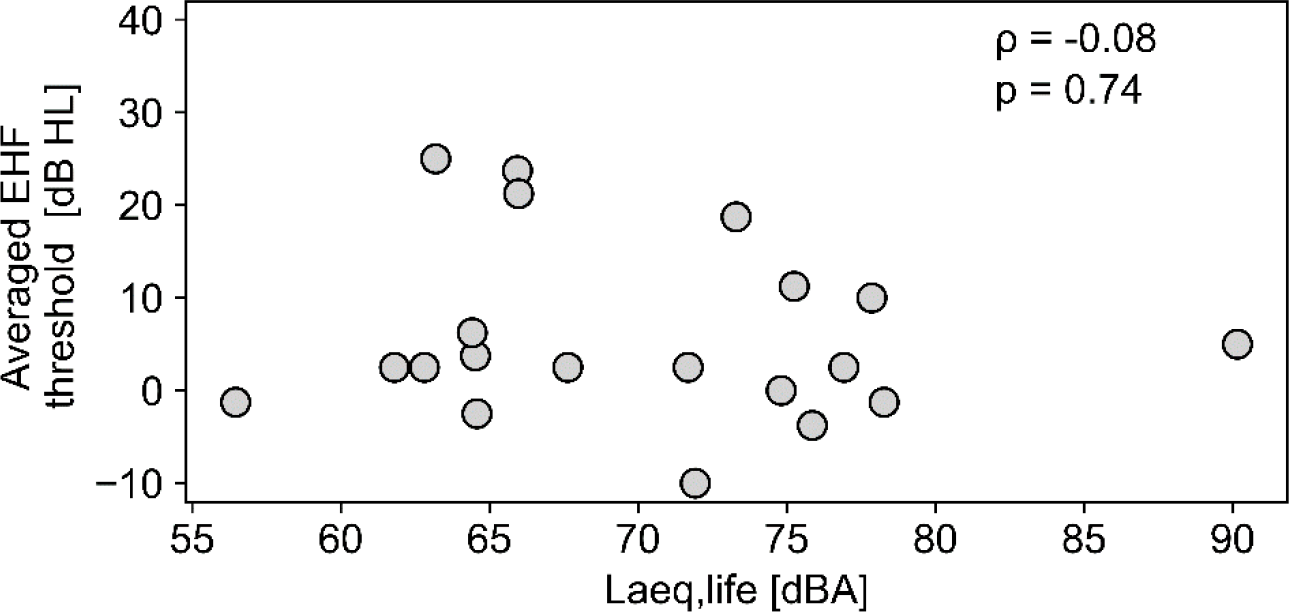
Spearman’s correlation analysis between Laeq,life and averaged EHF audiometric thresholds from 10 to 16 kHz.

**Figure 10.**
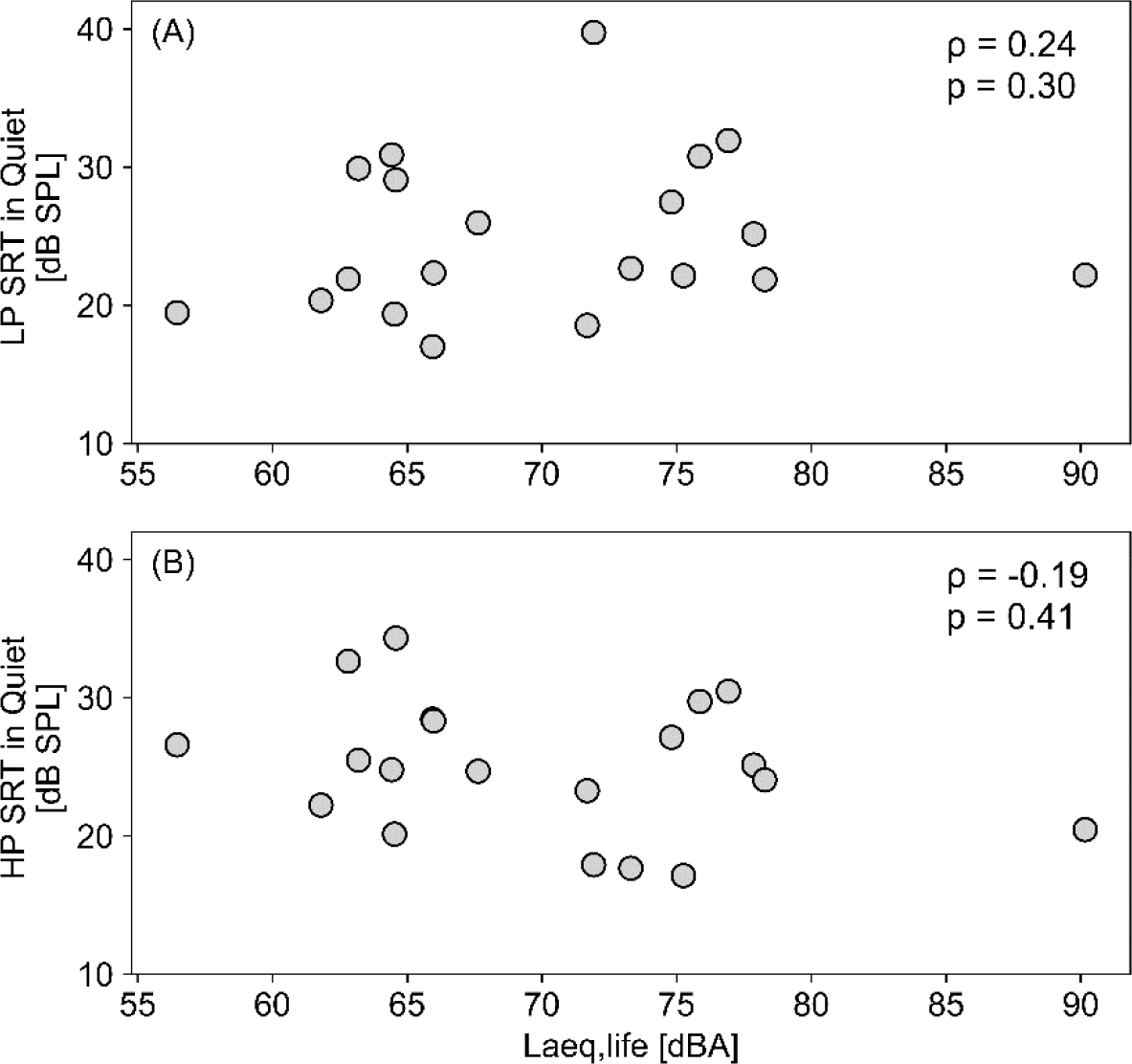
Spearman’s correlation analysis between Laeq,life and (A) LP SPiQ, and (B) HP SpiQ. Correlation coefficients and p-values are indicated in respective panels.

**Figure 11.**
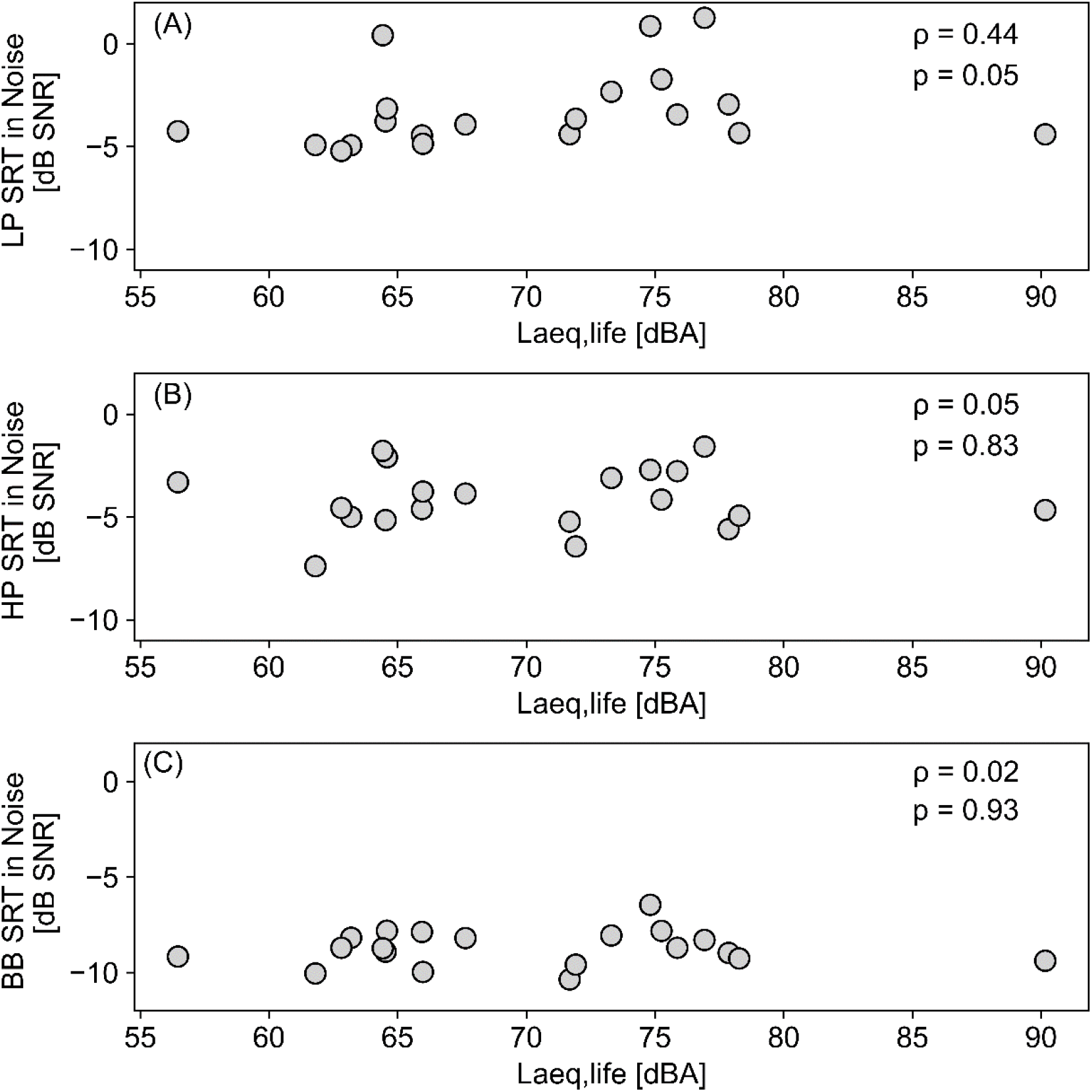
Spearman’s correlation analysis between Laeq,life and (A) LP SPiN, (B) HP SPiN and (C) BB SPiN reception thresholds. Correlation coefficients and p-values are indicated in respective panels.

**Figure 12.**
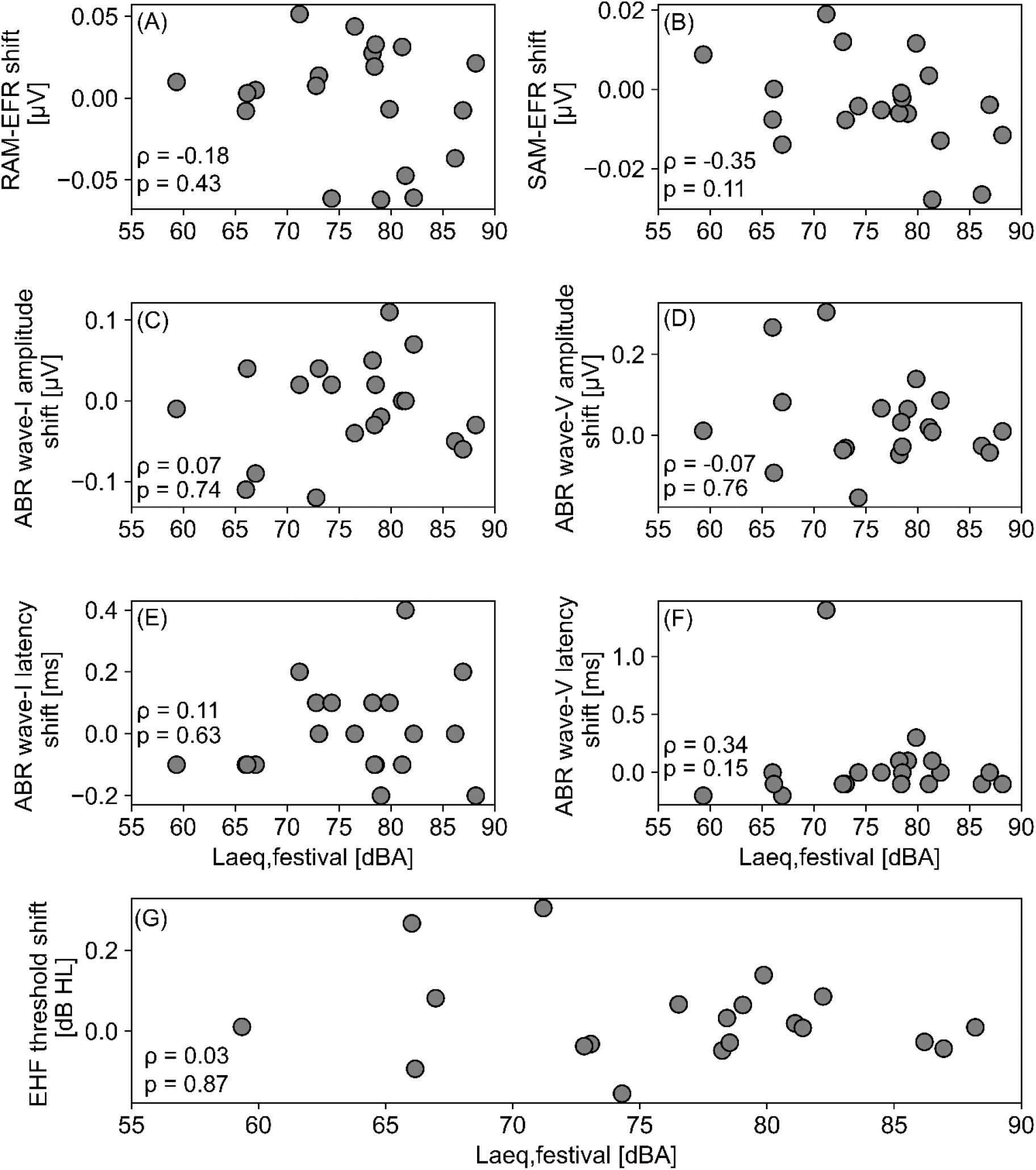
Spearman’s correlation between the Laeq,festival and the shift-amount of different markers from session-1 to session-2: (A) RAM-EFR magnitude, (B) SAM-EFR magnitude, (C) ABR wave-I amplitude, (D) ABR wave-V amplitude, (E) ABR wave-I latency, (F) ABR wave-V latency and (G) EHF threshold shifts. In each panel, a negative shift is an indicator of reduction in the corresponding from session-1 to session-2.

**Table 4.** U-values and significance levels of Mann Whitney U test for AEP measurements. Median test shifts from session 1 to session 2, minima and maxima are shown for subjects that have experienced Hearing Related Symptoms (HRS) and those who did not. Significant between-group differences are highlighted with *(p<0.05), **(p<0.01) or ***(p<0.001).

|  | Stimulus | no HRS (n=11) |  |  | HRS (n=9) |  |  | Mann-Whitney U |  |
| --- | --- | --- | --- | --- | --- | --- | --- | --- | --- |
|  |  | Median | Minimum | Maximum | Median | Minimum | Maximum | U | p |
| latency wave I | 11 Hz 80 dB | -0.10 | -0.20 | 0.10 | 0.05 | -0.10 | 0.40 | 78.00 | 0.03 * |
|  | 11 Hz 100 dB | -0.10 | -0.20 | 0.20 | 0.00 | 0.00 | 0.60 | 79.50 | 0.02 * |
|  | 120 Hz 80 dB | 0.00 | -0.30 | 0.20 | 0.15 | -0.70 | 1.20 | 66.00 | 0.23 |
|  | 120 Hz 100 dB | 0.10 | -0.30 | 1.20 | -0.05 | -0.70 | 0.20 | 27.50 | 0.10 |
|  | TB 1 kHz | 0.00 | -1.10 | 1.40 | -0.10 | -0.60 | 1.00 | 49.00 | 1.00 |
|  | TB 4 kHz | -0.20 | -1.00 | 1.90 | -0.15 | -1.00 | 0.80 | 39.50 | 0.66 |
| Latency wave III | 11 Hz 80 dB | 0.10 | -0.10 | 0.80 | 0.15 | -0.10 | 1.00 | 50.50 | 0.94 |
|  | 11 Hz 100 dB | 0.00 | -0.80 | 0.10 | 0.00 | -0.10 | 0.10 | 57.50 | 0.55 |
|  | 120 Hz 80 dB | 0.00 | -0.30 | 1.20 | -0.05 | -0.60 | 0.20 | 35.00 | 0.30 |
|  | 120 Hz 100 dB | 0.00 | -0.80 | 0.20 | 0.05 | -0.20 | 0.70 | 72.00 | 0.10 |
|  | TB 1 kHz | 0.10 | -0.70 | 1.50 | 0.20 | -1.20 | 0.70 | 48.00 | 0.94 |
|  | TB 4 kHz | 0.20 | -0.40 | 1.30 | 0.00 | -0.70 | 0.40 | 16.50 | 0.02 * |
| Latency wave V | 11 Hz 80 dB | -0.10 | -0.20 | 0.10 | 0.00 | -0.10 | 1.40 | 73.00 | 0.08 |
|  | 11 Hz 100 dB | 0.00 | -0.10 | 0.40 | 0.05 | -1.30 | 0.20 | 53.00 | 0.82 |
|  | 120 Hz 80 dB | 0.00 | -0.10 | 0.40 | -0.05 | -0.50 | 1.10 | 46.50 | 0.82 |
|  | 120 Hz 100 dB | -0.10 | -0.20 | 0.70 | 0.10 | -0.10 | 1.10 | 73.00 | 0.08 |
|  | TB 1 kHz | 0.10 | -0.40 | 1.40 | -0.05 | -0.60 | 0.10 | 29.00 | 0.13 |
|  | TB 4 kHz | 0.05 | -0.50 | 2.00 | -0.10 | -0.20 | 0.40 | 32.00 | 0.32 |
| Amplitude wave I | 11 Hz 80 dB | -0.01 | -0.12 | 0.07 | 0.01 | -0.05 | 0.11 | 52.00 | 0.88 |
|  | 11 Hz 100 dB | 0.01 | -0.15 | 0.13 | 0.03 | -0.20 | 0.20 | 56.00 | 0.66 |
|  | 120 Hz 80 dB | 0.02 | -0.08 | 0.17 | -0.00 | -0.18 | 0.17 | 45.00 | 0.77 |
|  | 120 Hz 100 dB | 0.02 | -0.08 | 0.10 | -0.02 | -0.13 | 0.08 | 31.00 | 0.18 |
|  | TB 1 kHz | -0.01 | -0.08 | 0.24 | -0.06 | -0.12 | 0.23 | 35.00 | 0.30 |
|  | TB 4 kHz | -0.01 | -0.07 | 0.04 | 0.06 | -0.05 | 0.16 | 71.00 | 0.04 * |
| Amplitude<br>wave III | 11 Hz<br>80 dB | 0.02 | -0.04 | 0.13 | 0.01 | -0.10 | 0.05 | 32.00 | 0.20 |
|  | 11 Hz<br>100 dB | -0.04 | -0.13 | 0.14 | -0.03 | -0.13 | 0.16 | 55.00 | 0.71 |
|  | 120 Hz<br>80 dB | -0.01 | -0.46 | 0.13 | -0.03 | -0.28 | 0.14 | 55.00 | 1.00 |
|  | 120 Hz<br>100 dB | -0.00 | -0.29 | 0.10 | -0.03 | -0.24 | 0.10 | 47.00 | 0.88 |
|  | TB<br>1 kHz | -0.03 | -0.17 | 0.08 | 0.03 | -0.04 | 0.24 | 71.00 | 0.11 |
|  | TB<br>4 kHz | -0.02 | -0.15 | 0.10 | 0.03 | -0.08 | 0.12 | 53.00 | 0.55 |
| Amplitude<br>wave V | 11 Hz<br>80 dB | 0.01 | -0.09 | 0.27 | 0.02 | -0.15 | 0.30 | 50.00 | 1.00 |
|  | 11 Hz<br>100 dB | 0.06 | -0.15 | 0.15 | 0.01 | -0.04 | 0.23 | 55.00 | 0.71 |
|  | 120 Hz<br>80 dB | 0.01 | -0.15 | 0.19 | 0.11 | -0.15 | 0.22 | 61.00 | 0.41 |
|  | 120 Hz<br>100 dB | 0.01 | -0.30 | 0.15 | 0.02 | -0.09 | 0.27 | 65.00 | 0.26 |
|  | TB<br>1 kHz | 0.01 | -0.08 | 0.06 | -0.00 | -0.08 | 0.03 | 45.00 | 0.77 |
|  | TB<br>4 kHz | 0.04 | -0.10 | 0.15 | -0.04 | -0.16 | 0.04 | 17.00 | 0.02 * |
| I/V<br>amplitude<br>ratio | 11 Hz<br>80 dB | -0.04 | -0.67 | 0.40 | -0.04 | -1.01 | 0.42 | 48.00 | 0.78 |
|  | 11 Hz<br>100 dB | 0.08 | -0.54 | 0.45 | 0.11 | -0.95 | 0.39 | 50.00 | 0.66 |
|  | 120 Hz<br>80 dB | 0.08 | -0.55 | 1.00 | -0.01 | -0.95 | 0.25 | 31.00 | 0.31 |
|  | 120 Hz<br>100 dB | 0.09 | -0.36 | 8.57 | -0.06 | -0.77 | 0.05 | 17.00 | 0.03 * |
|  | TB<br>1 kHz | -0.18 | -4.04 | 63.34 | -0.34 | -1.16 | 3.85 | 35.00 | 0.49 |
|  | TB<br>4 kHz | 0.07 | -0.45 | 16.34 | 0.50 | -0.21 | 13.93 | 50.00 | 0.41 |

## REFERENCES

Beach, E. F., Williams, W., & Gilliver, M. (2012). The objective-subjective assessment of noise: Young adults can estimate loudness of events and lifestyle noise. International Journal of Audiology, 51(6), 444–449. https://doi.org/10.3109/14992027.2012.658971

Bharadwaj, H. M., Masud, S., Mehraei, G., Verhulst, S., & Shinn-Cunningham, B. G. (2015). Individual differences reveal correlates of hidden hearing deficits. The Journal of neuroscience : the official journal of the Society for Neuroscience, 35(5), 2161–2172. https://doi.org/10.1523/JNEUROSCI.3915-14.2015

Bharadwaj, H. M., Verhulst, S., Shaheen, L., Liberman, M. C., & Shinn-Cunningham, B. G. (2014). Cochlear neuropathy and the coding of supra-threshold sound. Frontiers in systems neuroscience, 8, 26. https://doi.org/10.3389/fnsys.2014.00026

Bramhall, N., Beach, E. F., Epp, B., Le Prell, C. G., Lopez-Poveda, E. A., Plack, C. J., Schaette, R., Verhulst, S., & Canlon, B. (2019). The search for noise-induced cochlear synaptopathy in humans: Mission impossible? Hearing research, 377, 88–103. https://doi.org/10.1016/j.heares.2019.02.016

Bramhall, N. F., Konrad-Martin, D., & McMillan, G. P. (2018). Tinnitus and Auditory Perception After a History of Noise Exposure: Relationship to Auditory Brainstem Response Measures [Article]. Ear and Hearing, 39(5), 881–894. https://doi.org/10.1097/AUD.0000000000000544

Bramhall, N. F., Konrad-Martin, D., McMillan, G. P., & Griest, S. E. (2017). Auditory Brainstem Response Altered in Humans With Noise Exposure Despite Normal Outer Hair Cell Function [Article]. Ear and hearing, 38(1), e1–e12. http://www.embase.com/search/results?subaction=viewrecord&from=export&id=L620874933

Carter, L., Williams, W., Black, D., & Bundy, A. (2014). The leisure-noise dilemma: hearing loss or hearsay? What does the literature tell us? Ear and hearing, 35(5), 491–505. https://doi.org/10.1097/01.aud.0000451498.92871.20

Church, M. W., Zhang, J. S., Langford, M. M., & Perrine, S. A. (2013). ’Ecstasy’ enhances noise-induced hearing loss. Hearing research, 302, 96–106. https://doi.org/10.1016/j.heares.2013.05.007

Clark, W. W. (1991). Recent studies of temporary threshold shift (TTS) and permanent threshold shift (PTS) in animals. The Journal of the Acoustical Society of America, 90(1), 155–163. https://doi.org/10.1121/1.401309

Couth, S., Prendergast, G., Guest, H., Munro, K. J., Moore, D. R., Plack, C. J., Ginsborg, J., & Dawes, P. (2020). Investigating the effects of noise exposure on self-report, behavioral and electrophysiological indices of hearing damage in musicians with normal audiometric thresholds. Hearing research, 108021. https://doi.org/10.1016/j.heares.2020.108021

Degeest, S., Clays, E., Corthals, P., & Keppler, H. (2017). Epidemiology and Risk Factors for Leisure Noise-Induced Hearing Damage in Flemish Young Adults. Noise & health, 19(86), 10–19. https://doi.org/10.4103/1463-1741.199241

Degeest, S., Keppler, H., Corthals, P., & Clays, E. (2017). Epidemiology and risk factors for tinnitus after leisure noise exposure in Flemish young adults. International journal of audiology, 56(2), 121–129. https://doi.org/10.1080/14992027.2016.1236416

Derebery, M. J., Vermiglio, A., Berliner, K. I., Potthoff, M., & Holguin, K. (2012). Facing the music: pre- and postconcert assessment of hearing in teenagers. Otology & neurotology : official publication of the American Otological Society, American Neurotology Society [and] European Academy of Otology and Neurotology, 33(7), 1136–1141. https://doi.org/10.1097/MAO.0b013e31825f2328

Doğru, H., Tüz, M., & Uygur, K. (2003). Correlation between blood group and noise-induced hearing loss. Acta oto-laryngologica, 123(8), 941–942.

Emmerich, E., Richter, F., Hagner, H., Giessler, F., Gehrlein, S., & Dieroff, H. G. (2002). Effects of discotheque music on audiometric results and central acoustic evoked neuromagnetic responses. The international tinnitus journal, 8(1), 13–19.

Fernandez, K. A., Jeffers, P. W., Lall, K., Liberman, M. C., & Kujawa, S. G. (2015). Aging after noise exposure: acceleration of cochlear synaptopathy in “recovered” ears. The Journal of neuroscience : the official journal of the Society for Neuroscience, 35(19), 7509–7520. https://doi.org/10.1523/jneurosci.5138-14.2015

Ferrite, S., & Santana, V. (2005). Joint effects of smoking, noise exposure and age on hearing loss. Occupational medicine (Oxford, England), 55(1), 48–53. https://doi.org/10.1093/occmed/kqi002

Francart, T., van Wieringen, A., & Wouters, J. (2008). APEX 3: a multi-purpose test platform for auditory psychophysical experiments. Journal of neuroscience methods, 172(2), 283–293. https://doi.org/10.1016/j.jneumeth.2008.04.020

Fulbright, A. N. C., Le Prell, C. G., Griffiths, S. K., & Lobarinas, E. (2017). Effects of Recreational Noise on Threshold and Suprathreshold Measures of Auditory Function. Seminars in hearing, 38(4), 298–318. https://doi.org/10.1055/s-0037-1606325

Furman, A. C., Kujawa, S. G., & Liberman, M. C. (2013). Noise-induced cochlear neuropathy is selective for fibers with low spontaneous rates. Journal of neurophysiology, 110(3), 577–586. https://doi.org/10.1152/jn.00164.2013

Garrett, M., Vasilkov, V., Mauermann, M., Wilson, J. L., Henry, K. S., & Verhulst, S. (2020). Speech-in-noise intelligibility difficulties with age: the role of cochlear synaptopathy. bioRxiv, 2020.2006.2009.142950. https://doi.org/10.1101/2020.06.09.142950

Garrett, M., & Verhulst, S. (2019). Applicability of subcortical EEG metrics of synaptopathy to older listeners with impaired audiograms. Hearing research, 380, 150–165. https://doi.org/10.1016/j.heares.2019.07.001

Grinn, S. K., Wiseman, K. B., Baker, J. A., & Le Prell, C. G. (2017). Hidden Hearing Loss? No Effect of Common Recreational Noise Exposure on Cochlear Nerve Response Amplitude in Humans. Frontiers in neuroscience, 11, 465. https://doi.org/10.3389/fnins.2017.00465

Grose, J. H., Buss, E., & Elmore, H. (2019). Age-Related Changes in the Auditory Brainstem Response and Suprathreshold Processing of Temporal and Spectral Modulation. Trends in hearing, 23, 2331216519839615. https://doi.org/10.1177/2331216519839615

Grose, J. H., Buss, E., & Hall, J. W. (2017). Loud Music Exposure and Cochlear Synaptopathy in Young Adults: Isolated Auditory Brainstem Response Effects but No Perceptual Consequences [Article]. Trends in hearing, 21, 2331216517737417. https://doi.org/10.1177/2331216517737417

Guest, H., Munro, K. J., & Plack, C. J. (2017). Tinnitus with a normal audiogram: Role of high-frequency sensitivity and reanalysis of brainstem-response measures to avoid audiometric over-matching [Letter]. Hearing Research, 356, 116–117. https://doi.org/10.1016/j.heares.2017.10.002

Guest, H., Munro, K. J., Prendergast, G., Millman, R. E., & Plack, C. J. (2018). Impaired speech perception in noise with a normal audiogram: No evidence for cochlear synaptopathy and no relation to lifetime noise exposure [Article]. Hearing Research, 364, 142–151. https://doi.org/10.1016/j.heares.2018.03.008

Hakuba, N., Koga, K., Gyo, K., Usami, S. I., & Tanaka, K. (2000). Exacerbation of noise-induced hearing loss in mice lacking the glutamate transporter GLAST. The Journal of neuroscience : the official journal of the Society for Neuroscience, 20(23), 8750–8753. https://www.ncbi.nlm.nih.gov/pubmed/11102482

Hall, J. W. (2007). New Handbook of Auditory Evoked Responses. Pearson. https://books.google.be/books?id=DpZrQgAACAAJ

Henry, K. S., Kale, S., & Heinz, M. G. (2016). Distorted Tonotopic Coding of Temporal Envelope and Fine Structure with Noise-Induced Hearing Loss. The Journal of neuroscience : the official journal of the Society for Neuroscience, 36(7), 2227–2237. https://doi.org/10.1523/jneurosci.3944-15.2016

Hope, A. J., Luxon, L. M., & Bamiou, D. E. (2013). Effects of chronic noise exposure on speech-in-noise perception in the presence of normal audiometry. The Journal of laryngology and otology, 127(3), 233–238. https://doi.org/10.1017/S002221511200299X

Jensen, J. B., Lysaght, A. C., Liberman, M. C., Qvortrup, K., & Stankovic, K. M. (2015). Immediate and delayed cochlear neuropathy after noise exposure in pubescent mice [Article]. PLoS ONE, 10(5). https://doi.org/10.1371/journal.pone.0125160

Jokitulppo, J., Toivonen, M., & Bjork, E. (2006). Estimated leisure-time noise exposure, hearing thresholds, and hearing symptoms of Finnish conscripts. Military medicine, 171(2), 112–116. https://www.ncbi.nlm.nih.gov/pubmed/16578978

Keppler, H., Dhooge, I., Degeest, S., & Vinck, B. (2015). The effects of a hearing education program on recreational noise exposure, attitudes and beliefs toward noise, hearing loss, and hearing protector devices in young adults. Noise & health, 17(78), 253–262. https://doi.org/10.4103/1463-1741.165028

Keppler, H., Dhooge, I., & Vinck, B. (2015a). Hearing in young adults. Part I: The effects of attitudes and beliefs toward noise, hearing loss, and hearing protector devices. Noise & health, 17(78), 237–244. https://doi.org/10.4103/1463-1741.165024

Keppler, H., Dhooge, I., & Vinck, B. (2015b). Hearing in young adults. Part II: The effects of recreational noise exposure. Noise & health, 17(78), 245–252. https://doi.org/10.4103/1463-1741.165026

Keppler, H., & Dhooge, I. J. M. c. (2010). Optimization of the diagnosis of noise-induced hearing loss with otoacoustic emissions 2010.]. http://lib.ugent.be/catalog/rug01:001438088

Keshishzadeh, S., Garrett, M., Vasilkov, V., & Verhulst, S. (2020). The derived-band envelope following response and its sensitivity to sensorineural hearing deficits. Hearing Research, 392, 107979. https://doi.org/10.1016/j.heares.2020.107979

Keshishzadeh, S., Garrett, M., & Verhulst, S. (2021). Towards Personalized Auditory Models: Predicting Individual Sensorineural Hearing-Loss Profiles From Recorded Human Auditory Physiology. Trends in Hearing, 25, 2331216520988406.

Kolkhorst, F. W., Smaldino, J. J., Wolf, S. C., Battani, L. R., Plakke, B. L., Huddleston, S., & Hensley, L. D. (1998). Influence of fitness on susceptibility to noise-induced temporary threshold shift. Medicine and science in sports and exercise, 30(2), 289–293. https://www.ncbi.nlm.nih.gov/pubmed/9502359

Kollmeier, B., Warzybok, A., Hochmuth, S., Zokoll, M. A., Uslar, V., Brand, T., & Wagener, K. C. (2015). The multilingual matrix test: Principles, applications, and comparison across languages: A review. International journal of audiology, 54 *Suppl 2*, 3–16. https://doi.org/10.3109/14992027.2015.1020971

Konrad-Martin, D., Dille, M. F., McMillan, G., Griest, S., McDermott, D., Fausti, S. A., & Austin, D. F. (2012). Age-related changes in the auditory brainstem response. Journal of the American Academy of Audiology, 23(1), 18–35; quiz 74-15. https://doi.org/10.3766/jaaa.23.1.3

Kopke, R., Bielefeld, E., Liu, J., Zheng, J., Jackson, R., Henderson, D., & Coleman, J. K. (2005). Prevention of impulse noise-induced hearing loss with antioxidants. Acta oto-laryngologica, 125(3), 235–243. https://www.ncbi.nlm.nih.gov/pubmed/15966690

Kraaijenga, V. J. C., van Munster, J., & van Zanten, G. A. (2018). Association of Behavior With Noise-Induced Hearing Loss Among Attendees of an Outdoor Music Festival: A Secondary Analysis of a Randomized Clinical Trial. JAMA otolaryngology--head & neck surgery, 144(6), 490–497. https://doi.org/10.1001/jamaoto.2018.0272

Kujawa, S. G., & Liberman, M. C. (2009). Adding insult to injury: cochlear nerve degeneration after “temporary” noise-induced hearing loss. The Journal of neuroscience : the official journal of the Society for Neuroscience, 29(45), 14077–14085. https://doi.org/10.1523/JNEUROSCI.2845-09.2009

Kujawa, S. G., & Liberman, M. C. (2015). Synaptopathy in the noise-exposed and aging cochlea: Primary neural degeneration in acquired sensorineural hearing loss. Hearing research, 330(Pt B), 191–199. https://doi.org/10.1016/j.heares.2015.02.009

Kumar, U. A., Ameenudin, S., & Sangamanatha, A. V. (2012). Temporal and speech processing skills in normal hearing individuals exposed to occupational noise. Noise & health, 14(58), 100–105. https://doi.org/10.4103/1463-1741.97252

Le Prell, C. G. (2019). Effects of noise exposure on auditory brainstem response and speech-in-noise tasks: a review of the literature. International journal of audiology, 58(sup1), S3–S32. https://doi.org/10.1080/14992027.2018.1534010

Le Prell, C. G., Dell, S., Hensley, B., Hall, J. W., 3rd, Campbell, K. C. M., Antonelli, P. J., Green, G. E., Miller, J. M., & Guire, K. (2012). Digital music exposure reliably induces temporary threshold shift in normal-hearing human subjects. Ear and hearing, 33(6), e44-e58. https://doi.org/10.1097/AUD.0b013e31825f9d89

Liberman, M. C. (1982). Single-neuron labeling in the cat auditory nerve. Science (New York, N.Y.), 216(4551), 1239–1241. https://doi.org/10.1126/science.7079757

Liberman, M. C., Dodds, L. W., & Pierce, S. (1990). Afferent and efferent innervation of the cat cochlea: quantitative analysis with light and electron microscopy. The Journal of comparative neurology, 301(3), 443–460. https://doi.org/10.1002/cne.903010309

Liberman, M. C., Epstein, M. J., Cleveland, S. S., Wang, H., & Maison, S. F. (2016). Toward a differential diagnosis of hidden hearing loss in humans [Article]. PLoS ONE, 11(9). https://doi.org/10.1371/journal.pone.0162726

Liberman, M. C., & Kujawa, S. G. (2017). Cochlear synaptopathy in acquired sensorineural hearing loss: Manifestations and mechanisms [Review]. Hearing Research, 349, 138–147. https://doi.org/10.1016/j.heares.2017.01.003

Lin, H. W., Furman, A. C., Kujawa, S. G., & Liberman, M. C. (2011). Primary neural degeneration in the Guinea pig cochlea after reversible noise-induced threshold shift. Journal of the Association for Research in Otolaryngology : JARO, 12(5), 605–616. https://doi.org/10.1007/s10162-011-0277-0

Luts, H., Jansen, S., Dreschler, W., & Wouters, J. (2014). Development and normative data for the Flemish/Dutch Matrix test.

Makary, C. A., Shin, J., Kujawa, S. G., Liberman, M. C., & Merchant, S. N. (2011). Age-related primary cochlear neuronal degeneration in human temporal bones. Journal of the Association for Research in Otolaryngology : JARO, 12(6), 711–717. https://doi.org/10.1007/s10162-011-0283-2

Masterson, E. A., Themann, C. L., Luckhaupt, S. E., Li, J., & Calvert, G. M. (2016). Hearing difficulty and tinnitus among U.S. workers and non-workers in 2007. American journal of industrial medicine, 59(4), 290-300. https://doi.org/10.1002/ajim.22565

Mepani, A. M., Verhulst, S., Hancock, K. E., Garrett, M., Vasilkov, V., Bennett, K., de Gruttola, V., Liberman, M. C., & Maison, S. F. (2021). Envelope following responses predict speech-in-noise performance in normal-hearing listeners. Journal of Neurophysiology, 125(4), 1213–1222.

Mercier, V., Luy, D., & Hohmann, B. W. (2003). The sound exposure of the audience at a music festival. Noise & health, 5(19), 51–58. http://www.noiseandhealth.org/article.asp?issn=1463-1741;year = 2003;volume=5;issue=19;spage=51;epage=58;aulast=Mercier

Möhrle, D., Ni, K., Varakina, K., Bing, D., Lee, S. C., Zimmermann, U., Knipper, M., & Rüttiger, L. (2016). Loss of auditory sensitivity from inner hair cell synaptopathy can be centrally compensated in the young but not old brain. Neurobiology of Aging, 44, 173–184. https://doi.org/10.1016/j.neurobiolaging.2016.05.001

Opperman, D. A., Reifman, W., Schlauch, R., & Levine, S. (2006). Incidence of spontaneous hearing threshold shifts during modern concert performances. Otolaryngology--head and neck surgery : official journal of American Academy of Otolaryngology-Head and Neck Surgery, 134(4), 667–673. https://doi.org/10.1016/j.otohns.2005.11.039

OSHA, U. S. D. o. L.-O. S. a. H. A. (1974). https://www.osha.gov/laws-regs/regulations/standardnumber/1910/1910.95

Parthasarathy, A., Bartlett, E. L., & Kujawa, S. G. (2018). Age-related Changes in Neural Coding of Envelope Cues: Peripheral Declines and Central Compensation. Neuroscience. https://doi.org/10.1016/j.neuroscience.2018.12.007

Parthasarathy, A., & Kujawa, S. G. (2018). Synaptopathy in the aging cochlea: Characterizing early-neural deficits in auditory temporal envelope processing [Article]. Journal of Neuroscience, 38(32), 7108–7119. https://doi.org/10.1523/JNEUROSCI.3240-17.2018

Paul, B. T., Waheed, S., Bruce, I. C., & Roberts, L. E. (2017). Subcortical amplitude modulation encoding deficits suggest evidence of cochlear synaptopathy in normal-hearing 18-19 year olds with higher lifetime noise exposure. The Journal of the Acoustical Society of America, 142(5), El434. https://doi.org/10.1121/1.5009603

Petrescu, N. (2008). Loud music listening. McGill journal of medicine : MJM : an international forum for the advancement of medical sciences by students, 11(2), 169–176. https://www.ncbi.nlm.nih.gov/pubmed/19148318

Picton, T. W. (2011). Human Auditory Evoked Potentials. Plural Pub. https://books.google.be/books?id=z05QAQAAIAAJ

Pouryaghoub, G., Mehrdad, R., & Mohammadi, S. (2007). Interaction of smoking and occupational noise exposure on hearing loss: a cross-sectional study. BMC public health, 7, 137. https://doi.org/10.1186/1471-2458-7-137

Prendergast, G., Couth, S., Millman, R. E., Guest, H., Kluk, K., Munro, K. J., & Plack, C. J. (2019). Effects of Age and Noise Exposure on Proxy Measures of Cochlear Synaptopathy. Trends in hearing, 23, 2331216519877301. https://doi.org/10.1177/2331216519877301

Prendergast, G., Guest, H., Munro, K. J., Kluk, K., Leger, A., Hall, D. A., Heinz, M. G., & Plack, C. J. (2017). Effects of noise exposure on young adults with normal audiograms I: Electrophysiology. Hearing research, 344, 68–81. https://doi.org/10.1016/j.heares.2016.10.028

Prendergast, G., Millman, R. E., Guest, H., Munro, K. J., Kluk, K., Dewey, R. S., Hall, D. A., Heinz, M. G., & Plack, C. J. (2017). Effects of noise exposure on young adults with normal audiograms II: Behavioral measures [Article]. Hearing Research, 356, 74–86. https://doi.org/10.1016/j.heares.2017.10.007

Prendergast, G., Tu, W., Guest, H., Millman, R. E., Kluk, K., Couth, S., Munro, K. J., & Plack, C. J. (2018). Supra-threshold auditory brainstem response amplitudes in humans: Test-retest reliability, electrode montage and noise exposure [Article]. Hearing Research, 364, 38–47. https://doi.org/10.1016/j.heares.2018.04.002

Puel, J. L., Ruel, J., Gervais d’Aldin, C., & Pujol, R. (1998). Excitotoxicity and repair of cochlear synapses after noise-trauma induced hearing loss. Neuroreport, 9(9), 2109–2114. https://doi.org/10.1097/00001756-199806220-00037

Ramakers, G. G., Kraaijenga, V. J., Cattani, G., van Zanten, G. A., & Grolman, W. (2016). Effectiveness of Earplugs in Preventing Recreational Noise-Induced Hearing Loss: A Randomized Clinical Trial. JAMA otolaryngology--head & neck surgery, 142(6), 551–558. https://doi.org/10.1001/jamaoto.2016.0225

Ridley, C. L., Kopun, J. G., Neely, S. T., Gorga, M. P., & Rasetshwane, D. M. (2018). Using Thresholds in Noise to Identify Hidden Hearing Loss in Humans [Article]. Ear and Hearing, 39(5), 829–844. https://doi.org/10.1097/aud.0000000000000543

Ryan, A. F., Kujawa, S. G., Hammill, T., Le Prell, C., & Kil, J. (2016). Temporary and Permanent Noise-induced Threshold Shifts: A Review of Basic and Clinical Observations. Otology & neurotology : official publication of the American Otological Society, American Neurotology Society [and] European Academy of Otology and Neurotology, 37(8), e271–275. https://doi.org/10.1097/mao.0000000000001071

Ryberg, J. B. (2009). A national project to evaluate and reduce high sound pressure levels from music. Noise & health, 11(43), 124–128. https://doi.org/10.4103/1463-1741.50698

Safieddine, S., El-Amraoui, A., & Petit, C. (2012). The auditory hair cell ribbon synapse: from assembly to function. Annual review of neuroscience, 35, 509–528. https://doi.org/10.1146/annurev-neuro-061010-113705

Schaette, R., & McAlpine, D. (2011). Tinnitus with a normal audiogram: physiological evidence for hidden hearing loss and computational model. The Journal of neuroscience : the official journal of the Society for Neuroscience, 31(38), 13452–13457. https://doi.org/10.1523/JNEUROSCI.2156-11.2011

Schlauch, R. S., & Carney, E. (2012). The challenge of detecting minimal hearing loss in audiometric surveys. American journal of audiology, 21(1), 106–119. https://doi.org/10.1044/1059-0889(2012/11-0012)

Schmuzigert, N., Fostiropoulos, K., & Probst, R. (2006). Long-term assessment of auditory changes resulting from a single noise exposure associated with non-occupational activities. International journal of audiology, 45(1), 46–54. https://doi.org/10.1080/14992020500377089

Sergeyenko, Y., Lall, K., Charles Liberman, M., & Kujawa, S. G. (2013). Age-related cochlear synaptopathy: An early-onset contributor to auditory functional decline [Article]. Journal of Neuroscience, 33(34), 13686–13694. https://doi.org/10.1523/JNEUROSCI.1783-13.2013

Shaheen, L. A., Valero, M. D., & Liberman, M. C. (2015). Towards a Diagnosis of Cochlear Neuropathy with Envelope Following Responses. Journal of the Association for Research in Otolaryngology : JARO, 16(6), 727–745. https://doi.org/10.1007/s10162-015-0539-3

Skoe, E., & Tufts, J. (2018). Evidence of noise-induced subclinical hearing loss using auditory brainstem responses and objective measures of noise exposure in humans [Article]. Hearing Research, 361, 80–91. https://doi.org/10.1016/j.heares.2018.01.005

Smith, P. A., Davis, A., Ferguson, M., & Lutman, M. E. (2000). The prevalence and type of social noise exposure in young adults in England. Noise & health, 2(6), 41–56.

Smith, S. B., Krizman, J., Liu, C., White-Schwoch, T., Nicol, T., & Kraus, N. (2019). Investigating peripheral sources of speech-in-noise variability in listeners with normal audiograms. Hearing research, 371, 66–74. https://doi.org/10.1016/j.heares.2018.11.008

Spankovich, C., Bishop, C., Johnson, M. F., Elkins, A., Su, D., Lobarinas, E., & Le Prell, C. G. (2017). Relationship between dietary quality, tinnitus and hearing level: data from the national health and nutrition examination survey, 1999-2002. International journal of audiology, 56(10), 716- 722. https://doi.org/10.1080/14992027.2017.1331049

Spankovich, C., Le Prell, C. G., Lobarinas, E., & Hood, L. J. (2017). Noise History and Auditory Function in Young Adults With and Without Type 1 Diabetes Mellitus. Ear and hearing, 38(6), 724–735. https://doi.org/10.1097/AUD.0000000000000457

Stamper, G. C., & Johnson, T. A. (2015). Auditory function in normal-hearing, noise-exposed human ears. Ear and hearing, 36(2), 172–184. https://doi.org/10.1097/aud.0000000000000107

Svensson, E. B., Morata, T. C., Nylen, P., Krieg, E. F., & Johnson, A. C. (2004). Beliefs and attitudes among Swedish workers regarding the risk of hearing loss. International Journal of Audiology, 43(10), 585–593. https://doi.org/Doi 10.1080/14992020400050075

Tronstad, T. V., & Gelderblom, F. B. (2016). Sound exposure during outdoor music festivals. Noise & health, 18(83), 220.

Trzaskowski, B., Jędrzejczak, W. W., Piłka, E., Cieślicka, M., & Skarżyński, H. (2014). Otoacoustic emissions before and after listening to music on a personal player. Medical science monitor : international medical journal of experimental and clinical research, 20, 1426–1431. https://doi.org/10.12659/MSM.890747

Upile, T., Sipaul, F., Jerjes, W., Singh, S., Nouraei, R., Maaytah, M., Andrews, P., Graham, J., Hopper, C., & Wright, A. (2007). The acute effects of alcohol on auditory thresholds. BMC ear, nose, and throat disorders, 7, 4. https://doi.org/10.1186/1472-6815-7-4

Valderrama, J. T., Alvarez, I., de la Torre, A., Segura, J. C., Sainz, M., & Vargas, J. L. (2012). Recording of auditory brainstem response at high stimulation rates using randomized stimulation and averaging. Journal of the Acoustical Society of America, 132(6), 3856–3865. https://doi.org/10.1121/1.4764511

Valderrama, J. T., Beach, E. F., Yeend, I., Sharma, M., Van Dun, B., & Dillon, H. (2018). Effects of lifetime noise exposure on the middle-age human auditory brainstem response, tinnitus and speech-in-noise intelligibility. Hearing research, 365, 36–48. https://doi.org/10.1016/j.heares.2018.06.003

Vasilkov, V., Garrett, M., Mauermann, M., & Verhulst, S. (2021). Enhancing the sensitivity of the envelope-following response for cochlear synaptopathy screening in humans: The role of stimulus envelope. Hear Res, 400, 108132. https://doi.org/10.1016/j.heares.2020.108132

Vasilkov, V., & Verhulst, S. (2019). Towards a differential diagnosis of cochlear synaptopathy and outer-hair-cell deficits in mixed sensorineural hearing loss pathologies. medRxiv, 19008680. https://doi.org/10.1101/19008680

Verhulst, S., Altoe, A., & Vasilkov, V. (2018). Computational modeling of the human auditory periphery: Auditory-nerve responses, evoked potentials and hearing loss. Hearing research, 360, 55–75. https://doi.org/10.1016/j.heares.2017.12.018

Viana, L. M., O’Malley, J. T., Burgess, B. J., Jones, D. D., Oliveira, C. A. C. P., Santos, F., Merchant, S. N., Liberman, L. D., & Liberman, M. C. (2015). Cochlear neuropathy in human presbycusis: Confocal analysis of hidden hearing loss in post-mortem tissue [Article]. Hearing Research, 327, 78–88. https://doi.org/10.1016/j.heares.2015.04.014

Wang, Y., Hirose, K., & Liberman, M. C. (2002). Dynamics of noise-induced cellular injury and repair in the mouse cochlea. Journal of the Association for Research in Otolaryngology : JARO, 3(3), 248–268. https://doi.org/10.1007/s101620020028

Wei, W., Heinze, S., Gerstner, D. G., Walser, S. M., Twardella, D., Reiter, C., Weilnhammer, V., Perez-Alvarez, C., Steffens, T., & Herr, C. E. W. (2017). Audiometric notch and extended high-frequency hearing threshold shift in relation to total leisure noise exposure: An exploratory analysis. Noise & health, 19(91), 263–269. https://doi.org/10.4103/nah.NAH_28_17

Widen, S. E., Holmes, A. E., & Erlandsson, S. I. (2006). Reported hearing protection use in young adults from Sweden and the USA: effects of attitude and gender. International journal of audiology, 45(5), 273–280. https://doi.org/10.1080/14992020500485676

Williams, W., Carter, L., & Seeto, M. (2015). Pure tone hearing thresholds and leisure noise: Is there a relationship? Noise & health, 17(78), 358–363. https://doi.org/10.4103/1463-1741.165066

Yamane, H., Nakai, Y., Takayama, M., Iguchi, H., Nakagawa, T., & Kojima, A. (1995). Appearance of free radicals in the guinea pig inner ear after noise-induced acoustic trauma. European archives of oto-rhino-laryngology : official journal of the European Federation of Oto-Rhino-Laryngological Societies (EUFOS) : affiliated with the German Society for Oto-Rhino-Laryngology - Head and Neck Surgery, 252(8), 504–508. https://doi.org/10.1007/bf02114761

Yassi, A., Pollock, N., Tran, N., & Cheang, M. (1993). Risks to hearing from a rock concert. Canadian family physician Medecin de famille canadien, 39, 1045–1050. https://www.ncbi.nlm.nih.gov/pubmed/8499785 https://www.ncbi.nlm.nih.gov/pmc/PMC2379666/

Yeend, I., Beach, E. F., Sharma, M., & Dillon, H. (2017). The effects of noise exposure and musical training on suprathreshold auditory processing and speech perception in noise [Article]. Hearing Research, 353, 224–236. https://doi.org/10.1016/j.heares.2017.07.006

